# The first Miocene fossils from coastal woodlands in the southern East African Rift

**DOI:** 10.1101/2021.12.16.472914

**Authors:** René Bobe, Vera Aldeias, Zeresenay Alemseged, Will Archer, Georges Aumaître, Marion K. Bamford, Dora Biro, Didier L. Bourlès, David R. Braun, Cristian Capelli, João d’Oliveira Coelho, Jörg M. Habermann, Jason J. Head, Karim Keddadouche, Kornelius Kupczik, Anne-Elisabeth Lebatard, Tina Lüdecke, Amélia Macôa, Felipe I. Martínez, Jacinto Mathe, Clara Mendes, Luis Meira Paulo, Maria Pinto, Thomas A. Püschel, Frederico Tátá Regala, Mark Sier, Maria Joana Ferreira da Silva, Marc Stalmans, Susana Carvalho

**Author notes:** Corresponding author René Bobe.

## Abstract

The Miocene is a key time in the evolution of African mammals and their ecosystems witnessing the origin of the African apes and the isolation of eastern coastal forests through an expanding biogeographic arid corridor. Until recently, however, Miocene sites from the southeastern regions of the continent were unknown. Here we report discovery of the first Miocene fossil teeth from the shoulders of the Urema Rift in Gorongosa National Park, Mozambique, at the southern East African Rift System. We provide the first 1) radiometric age determinations of the fossiliferous Mazamba Formation, 2) reconstructions of past vegetation in the region based on pedogenic carbonates and fossil wood, and 3) description of fossil teeth from the southern rift. Gorongosa is unique in the East African Rift System in combining marine invertebrates, marine vertebrates, terrestrial mammals, and fossil woods in coastal paleoenvironments. The Gorongosa fossil sites offer the first evidence of persistent woodlands and forests on the coastal margins of southeastern Africa during the Miocene, and an exceptional assemblage of fossil vertebrates including new species. Further work will allow the testing of hypotheses positing the formation of a northeast-southwest arid corridor isolating species on the eastern coastal forests from those elsewhere in Africa.

**Brief:** The Miocene is a key time in the evolution of African mammals and their ecosystems encompassing hominine origins and the establishment of an arid corridor that isolated eastern Africa’s coastal forests. Until now, however, Miocene sites from southeastern Africa have been unknown. We report the discovery of the first Miocene fossil sites from Gorongosa National Park, Mozambique, and show that these sites formed in coastal settings. We provide radiometric ages for the fossiliferous sediments, reconstructions of past vegetation based on stable isotopes and fossil wood, and a description of the first fossil teeth from the region. Gorongosa is the only paleontological site in the East African Rift that combines fossil woods, marine invertebrates, marine vertebrates, and terrestrial mammals. Gorongosa offers the first evidence of persistent woodlands and forests on the coastal margins of southeastern Africa during the Miocene.

## Introduction

Much of our knowledge about African Miocene vertebrates and their environments derives from paleontological sites along the East African Rift System (EARS). However, considerable geographic and temporal gaps in the fossil record obscure a full appreciation of past biodiversity, biogeography, and ecosystem evolution on the continent. For example, until recently there were no sites with Miocene mammals in the southern 1,500 km of the EARS (Figure 1). Thus, the Miocene faunas and ecosystems of this southern region have remained virtually unknown. Furthermore, none of the well-known Miocene fossil sites in the EARS provides evidence of eastern African coastal forests, a major ecosystem that may have played a key role in hominin origins and the evolution of several mammalian lineages (Joordens, Feibel, Vonhof, Schulp, & Kroon, 2019; Kingdon, 2003). Although the necessity of documenting new fossil sites in previously unknown areas is widely appreciated and advocated (Almécija et al., 2021; Cote, 2018), discovering entirely new paleontological areas is a rare event (d’Oliveira Coelho, Anemone, & Carvalho, 2021). Here we describe the first dentognathic specimens of fossil vertebrates discovered in the East African Rift of central Mozambique. The specimens derive from the Mazamba Formation on the eastern shoulder of the Urema Rift in Gorongosa National Park (GNP) (Figure 2) (Habermann et al., 2019).

**Figure 1.**
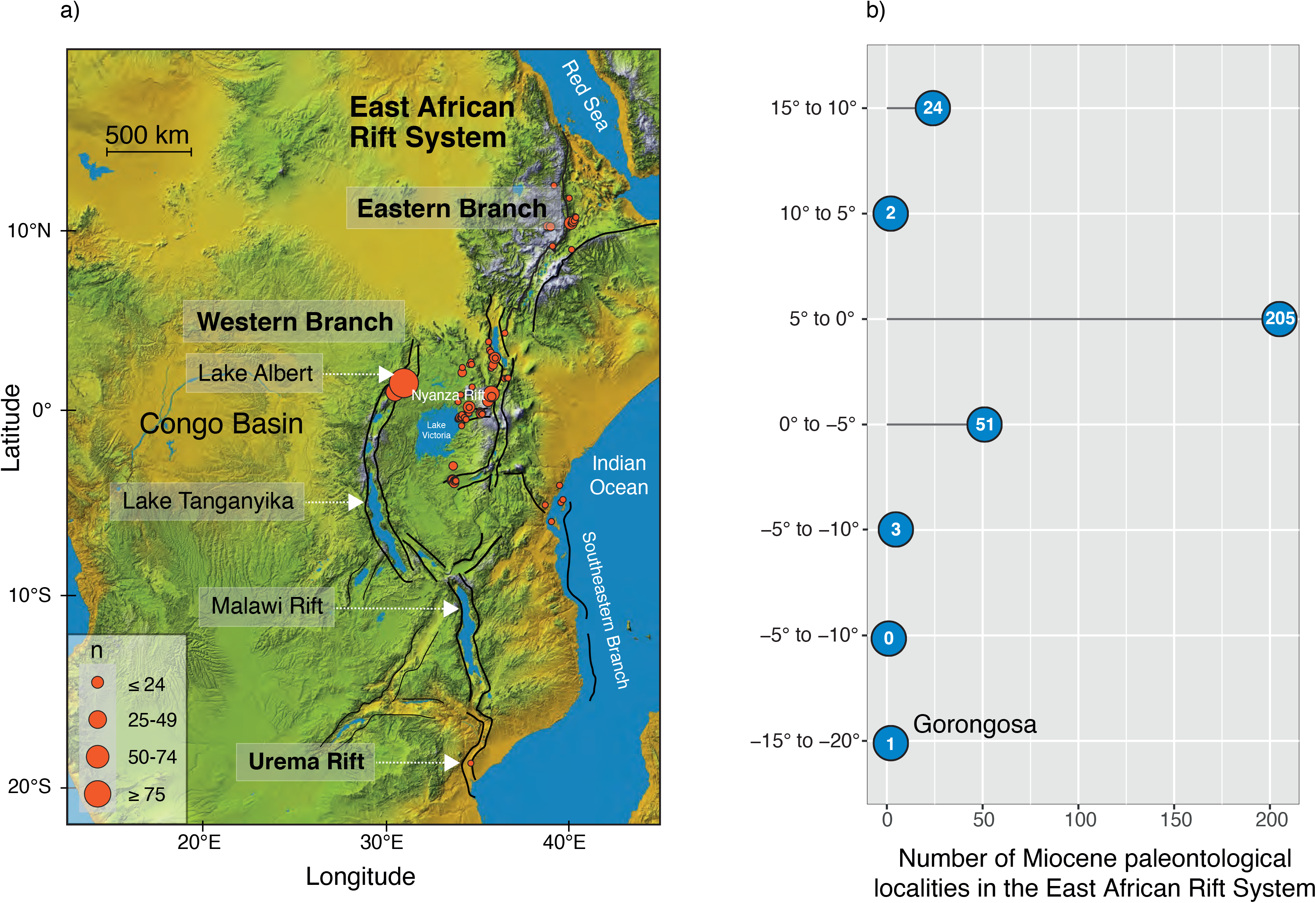
**A)** The East African Rift System (EARS) with the Eastern Branch, the Western Branch, and some of the major basins and rifts, including the Urema Graben at its southern end. The development of the EARS since the Miocene has played a major role in shaping the physical environments and modifying the conditions under which plants and animals have been evolving in eastern Africa. Base map from Nasa Shuttle Radar Topography Mission (https://www2.jpl.nasa.gov/srtm/). **B)** Number of Miocene paleontological localities along the EARS by latitude. There are many Miocene localities in the rift near the equator, but the record away from the equator, especially to the south, is very sparse. Gorongosa is the only Miocene paleontological locality in the southern ∼1500 km of the EARS. Locality data from the Paleobiology Database https://paleobiodb.org/classic.

**Figure 2.**
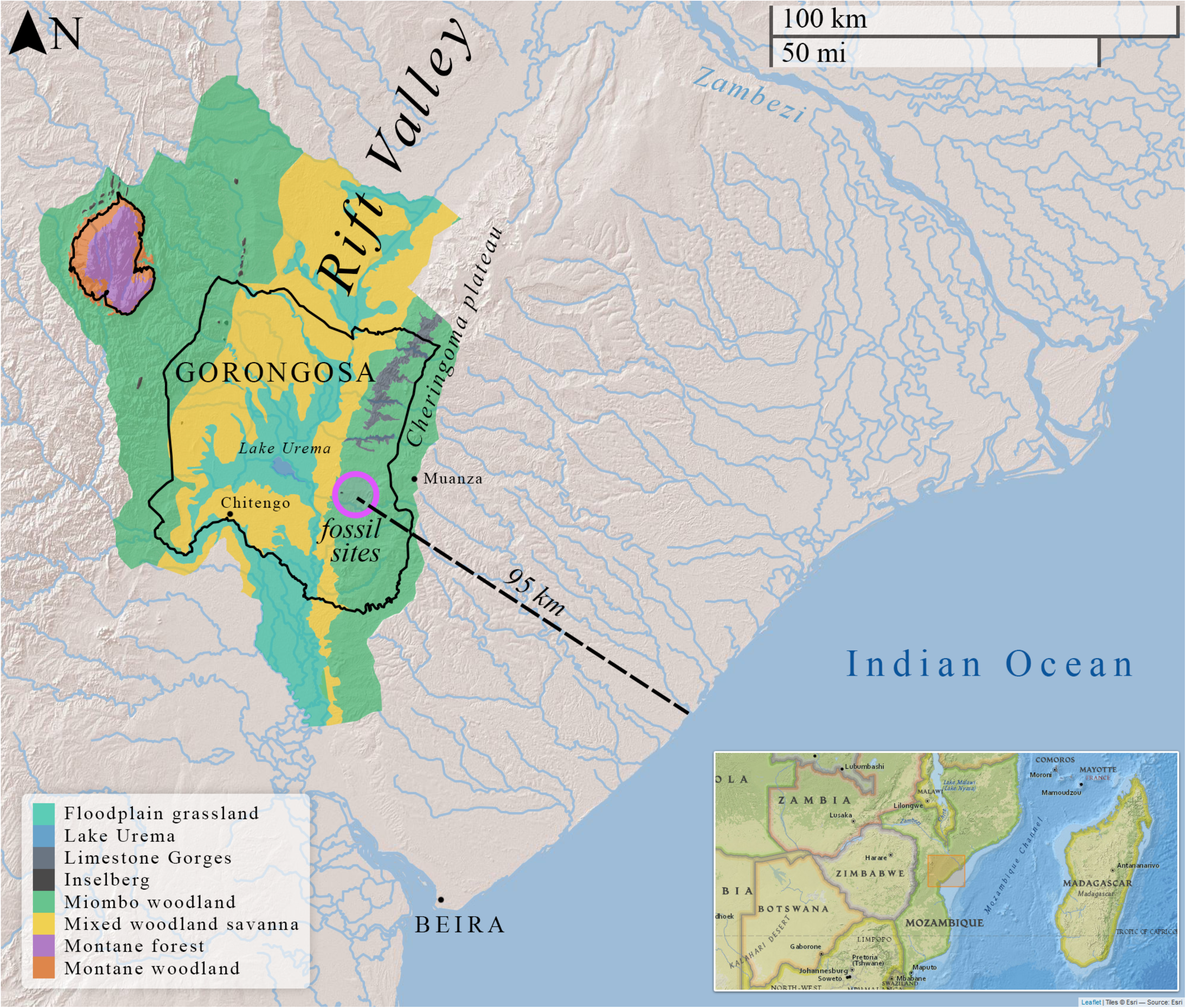
Map of Gorongosa National Park along the East African Rift Valley, with the Cheringoma Plateau to the east and Mount Gorongosa to the northwest. The park hosts a wide range of environments. The new paleontological sites on the Cheringoma Plateau are ∼95 km from the coast.

Cosmogenic nuclide dating presented here indicates that the Gorongosa paleontological localities are of Miocene age. These localities formed under estuarine conditions and represent the first samples of eastern African coastal woodlands and forests in the Miocene. The emerging fossil record from Gorongosa opens the possibility of testing, for the first time, key hypotheses about an expanding northeast-southwest arid corridor that would have isolated the eastern coastal forests from those in the central parts of Africa, and for exploring the importance of these processes for hominin origins (Kingdon, 2003). Gorongosa Park is now well known for its successful wildlife restoration project (Bouley, Paulo, Angela, Du Plessis, & Marneweck, 2021; Pringle, 2017; Stalmans, Massad, Peel, Tarnita, & Pringle, 2019), and these new paleontological sites in the park open a unique window on the fauna and environments of ancient Africa.

### Brief history of vertebrate paleontological research in the Cenozoic of Mozambique

Until recently, the fossil record of Cenozoic vertebrates in Mozambique has been sparse. More than a century and a half ago, J. Kirk reported on fossil finds of mammals and reptiles along the lower Zambezi River, but these were mixed with pottery and were likely to be recent, as they represented extant species (Kirk, 1864). In 1977, J. Harris briefly described a left upper premolar (P4) of the proboscidean *Deinotherium* found in coastal beach sands near Praia de Morrungulo, east of Massinga in Inhambane Province (Harris, 1977). This specimen is comparable in morphology and dimensions to Plio-Pleistocene *Deinotherium bozasi*, but the original provenance and geological context of the fossil is unknown. In 1998, a European speleological team surveyed the Cheringoma Plateau’s karstic system in Sofala Province (Figure 2), and the results published by M. Laumanns (2001) describe a series of caves near and within GNP. Some of the caves were indicated to have sedimentary infilling and archaeological potential, but there was no mention of any fossil vertebrates. Based on these reports, M. Pickford undertook brief surveys south of Morrungulo in 2012, and of GNP in 2012 and 2013. He discovered well-preserved fossil wood at Menguere Hill (=Mhengere in Pickford’s reports) (Figure 3) and found postcranial fragments of fossil mammals of uncertain taxonomic affinities north of the Muaredzi River (Pickford, 2012, 2013). According to Pickford, “The fossils are not well-preserved, and no teeth were found. Comparison with a wide range of mammals failed to provide definite identifications…” (Pickford, 2013: 8). At this time, J. Mercader and P. Sillén undertook archeological surveys of the Gorongosa region including some caves described by Laumanns (2001). They reported Holocene bones and teeth as well as Middle and Later Stone Age artifacts (Mercader & Sillén, 2013), but the bones illustrated by Mercader and Sillén are modern warthog, *Phacochoerus africanus*, and modern bush pig, *Potamochoerus larvatus*.

**Figure 3.**
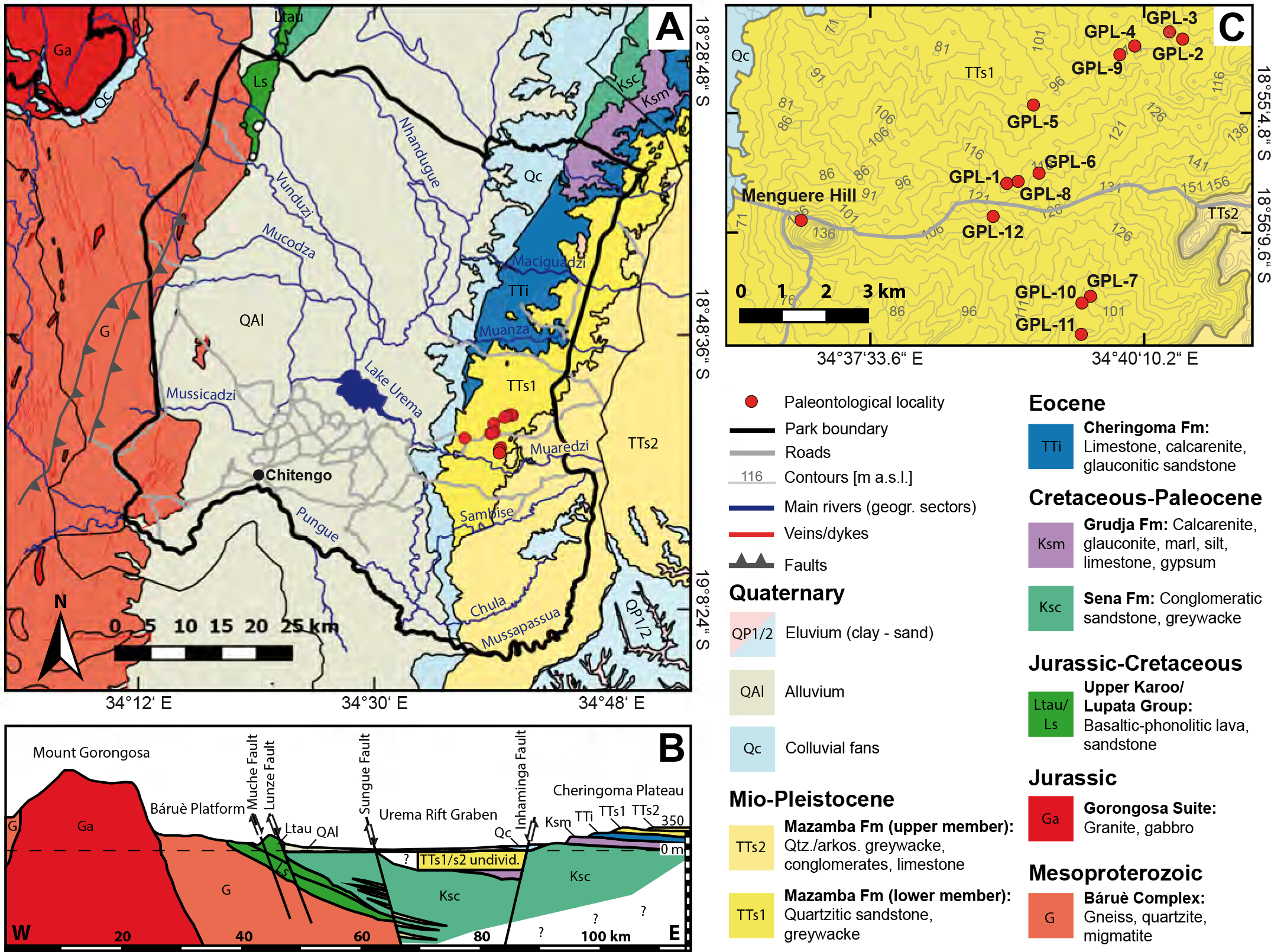
Gorongosa Paleontological Localities and geological formations. **A)** Geological map of Gorongosa National Park and surrounding areas. **B)** Vertical geological cross-section of the Urema Rift stretching from Mount Gorongosa to Inhaminga village. **C)** Map section showing the locations of the fossiliferous sites (GPL = Gorongosa Paleontological Locality). Figure modified from Habermann et al. (2019) and references therein, with new paleontological localities added.

In 2014, Gorongosa National Park authorities invited S. Carvalho to form a long-term paleontological, archaeological, speleological, and primatological research project. Carvalho created the Paleo-Primate Project Gorongosa (PPPG), an international and interdisciplinary endeavor combining paleontological and primatological approaches aiming to understand the deep time evolutionary history of the region. A central goal of PPPG is the mentoring and training of Mozambican students in areas of paleontology, archaeology, and primatology. In 2016, PPPG initiated systematic surveys of potentially fossiliferous sites within GNP, both in open-air sedimentary exposures of the Mazamba Formation and in the karstic cave systems of the Cheringoma Plateau. PPPG surveyed the vertebrate localities identified by Pickford in 2013 and these yielded several fragmentary fossils. After extensive surveys in new areas during the 2016 field season, the team discovered the first localities with teeth of fossil mammals the southern rift. After four field seasons (2016-2019), PPPG had documented hundreds of fossil vertebrates and fossil wood from 11 paleontological localities (Figure 3, Supporting Information Figure S1, Table 1). Some of these specimens are described below.

**Table 1.**
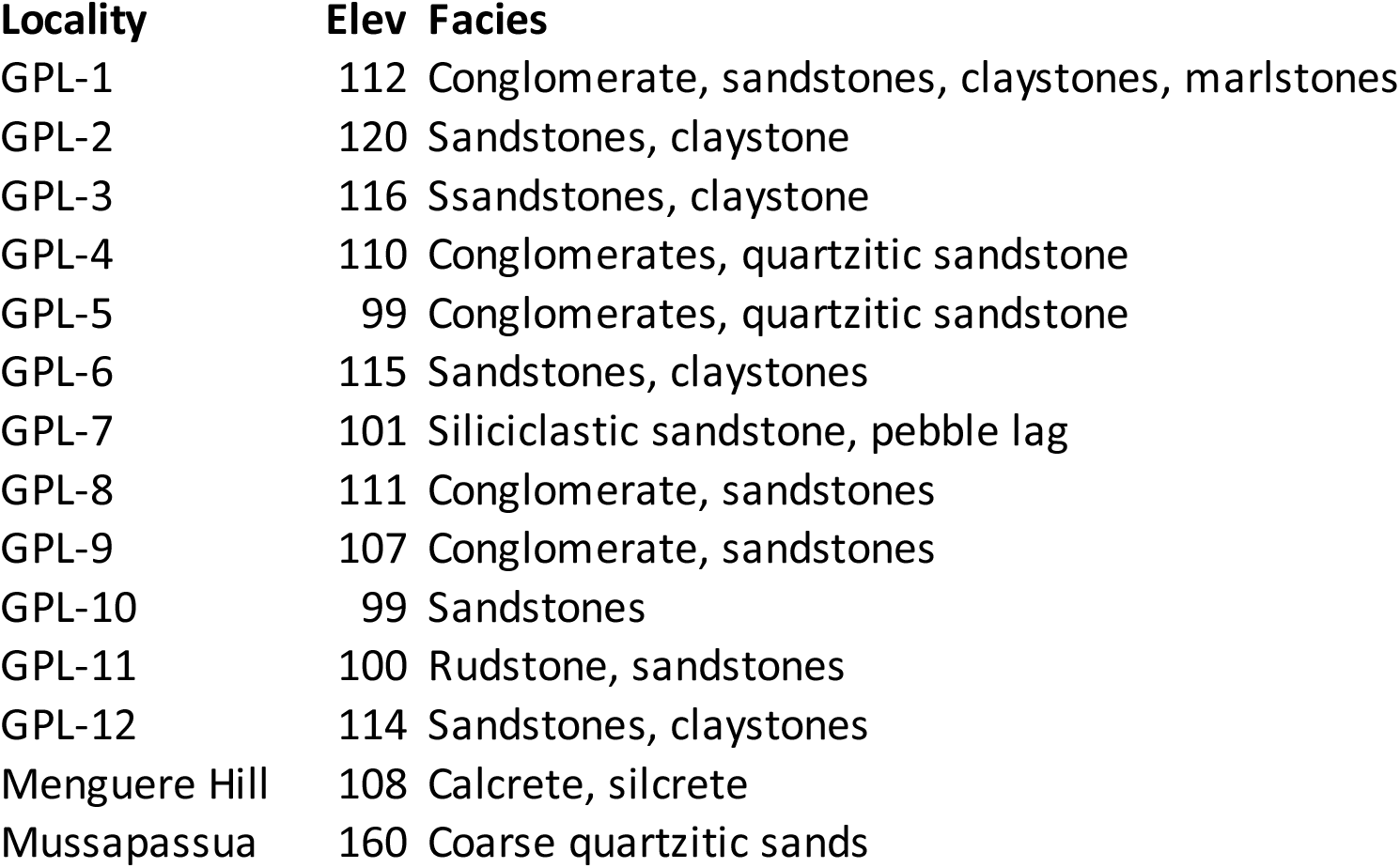
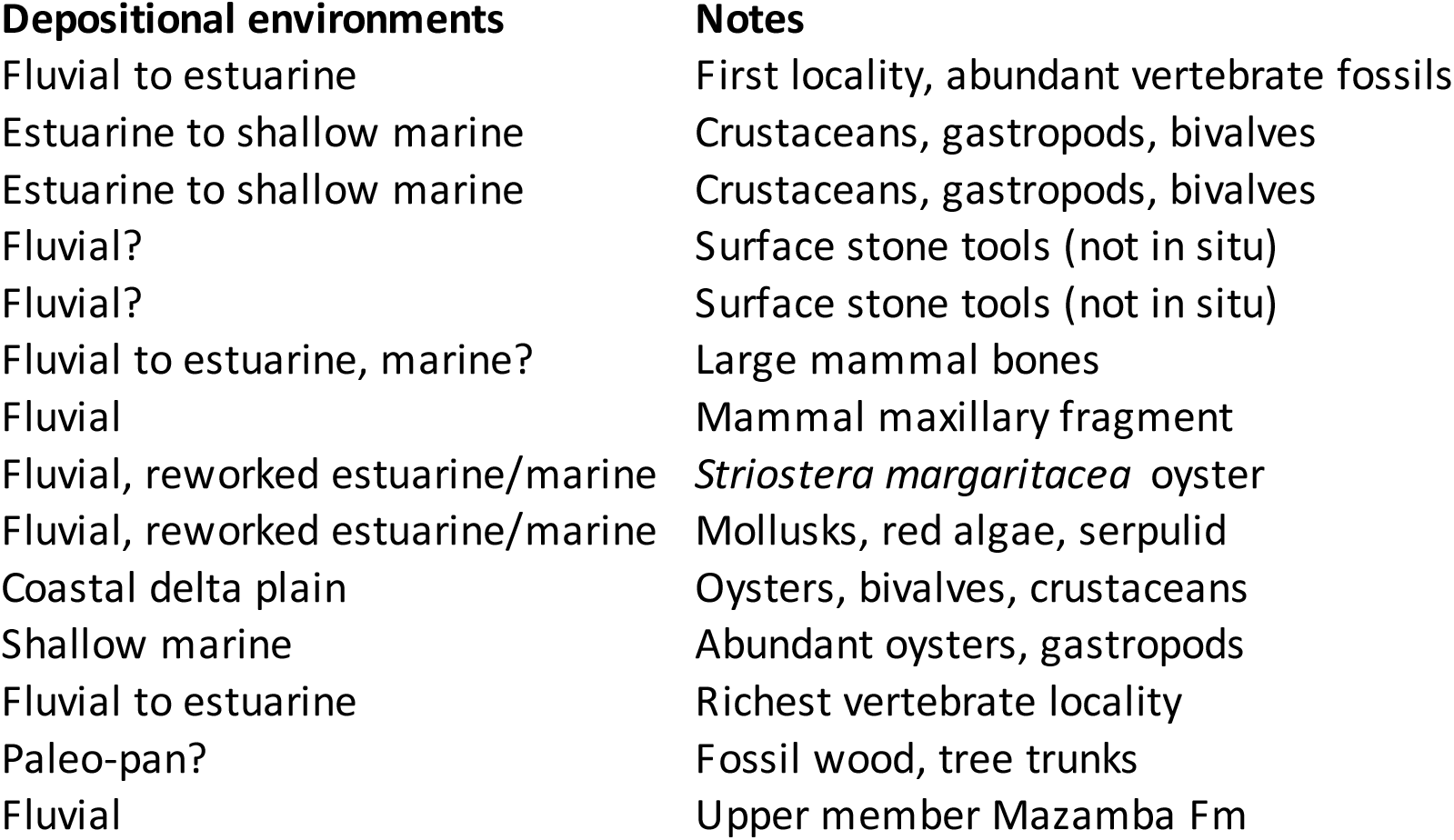
Gorongosa paleontological localities and depositional environments.

### The Urema Graben and the Cheringoma Plateau

At the southern end of the EARS, the Urema Graben crosses GNP along an approximately north-south axis, with the Cheringoma Plateau on the east and Mount Gorongosa dominating the northwestern region (Figures 2 and 3). Mount Gorongosa was formed by a granitic intrusion during the Jurassic (Real, 1966), but the Urema Graben represents one of the youngest sections of the EARS and may have been formed as recently as the Plio-Pleistocene (Grantham, Marques, Wilson, Manhiça, & Hartzer, 2011; Macgregor, 2015). The eastern shoulder of the Urema Graben is the Cheringoma Horst, an uplifted block bounded by the Inhaminga Fault on the west between the Pungue and Zambezi Rivers (Flores, 1973). Several geological formations are exposed in the Cheringoma Plateau, including the Sena Formation (Cretaceous), the Grudja Formation (with late Cretaceous and early Tertiary levels), the Cheringoma Formation (Eocene nummulitic limestones), and the Mazamba Formation (Mazamba sands attributed to the Miocene) (Flores, 1973; GTK_Consortium, 2006; Real, 1966) (Figure 3). Karstification of the Eocene limestones produced the extensive cave system of the Cheringoma Plateau (Laumanns, 2001).

Previous geological surveys of the region have determined that Cenozoic marine deposits begin with the glauconitic sands of the Grudja Formation, with one sample yielding a potassium-argon date of 60 Ma (Flores, 1973). Eocene marine deposits continue with the Cheringoma Formation, a limestone sequence with a thickness of 70-80 m containing abundant nummulites, echinoids, bryozoans, and gastropods deposited in warm, shallow (neritic) waters. The marine invertebrates (e.g., species of the annelid *Tubulostium*) indicate a middle to upper Eocene age for the Cheringoma Formation (Grantham et al., 2011; GTK_Consortium, 2006; Lächelt, 2004; Real, 1966; Teale, 1923). These Paleocene and Eocene deposits of the Grudja and Cheringoma formations represent a marine transgression with shorelines migrating westward (Flores, 1973).

The sedimentary deposits overlying the Cheringoma Formation constitute the Mazamba Formation, named after exposures along the Mazamba River 25 km southwest of Inhaminga in the Cheringoma Plateau. At the type locality in the upper Mazamba River this formation attains 140 m in thickness (Flores, 1973; Real, 1966). These deposits are separated from the underlying Cheringoma Formation by a well-defined erosional unconformity resulting from marine regression (Flores, 1973; Grantham et al., 2011). According to Flores (1973: 105), “There is an erosional unconformity between the Eocene and the Miocene, with no intervening Oligocene, indicating considerable uplift in post-Eocene-pre-Miocene times”. In the 1968 geological map of Mozambique, the Mazamba Formation is divided into two members separated by a chert horizon (as reproduced in Tinley 1977). The lower member (“grés de cor púrpura”, or purple clays/sands) (TT_S1_ in the 1968 geological map; Figure 3) is composed of purplish to reddish medium-grained argillaceous sands, which contain gastropods, bivalves, crustaceans, and foraminifera, and are interpreted to be littoral marine intercalated with deltaic deposits (Real, 1966). The upper member (TT_S2_) is referred to as the Inhaminga beds (“camadas de Inhaminga”), composed of medium to coarse arkosic sands with some irregular conglomerate layers (Figure 3). Real (1966) considers the lower member to be of Miocene age and the upper member to be of late Miocene to Pliocene age, but the criteria used to reach this conclusion are not specified. Lächelt (2004: 146-155) states that “The Mazamba Formation, which consists of conglomerates and sands, is the most distinctive Miocene formation” and attributes the sequence to the lower Miocene, but then he states that the Mazamba Formation correlates laterally with the Inhaminga Purple Sands. Although there are some discrepancies and contradictions (see Supporting Information: Geological Issues), most previous descriptions focused on the geology of the Cheringoma region consider the lower part of the Mazamba Formation to be of Miocene age and the upper part of the sequence to extend into the Pliocene (Kristina Arvidsson, 2010; K. Arvidsson et al., 2011; Habermann et al., 2019; Laumanns, 2001; Pickford, 2013; Tinley, 1977). Thus, we use the term Mazamba Formation to refer to the Mazamba/Inhaminga sequence in the Cheringoma Horst, with two informal members, a lower member and an upper member separated by a chert horizon. In the field, we identified the nodular chert layer separating the lower and upper sequences and undertook geological and paleontological surveys of both lower and upper deposits.

The dating of these sedimentary sequences has been hampered by the lack of radio-isotopic age determinations. Neogene volcanism has been less intensively developed in the southern EARS than in regions to the north (e.g., Afar, Main Ethiopian Rift, Omo-Turkana Basin, Kenya Rift), and volcanic ash layers amenable to radiometric dating seem to be rare. A basaltic intrusion ∼8 km northwest of the Cundué River cuts through the Grudja and Cheringoma Formations, and thus would be younger than the Eocene (Real, 1966). This basalt, however, has not been radiometrically dated. In a regional context, recent research on the Zambezi Delta by Ponte and colleagues has identified a major unconformity at the end of the Oligocene related to uplift of the South African Plateau (Baby, Guillocheau, Boulogne, Robin, & Dall’Asta, 2018), with the ‘Mazamba sands’ deposited above this unconformity during the early Miocene (Aquitanian and Burdigalian stages) (Ponte et al., 2019). See Supporting Information: Geological Issues for further details.

## Materials and Methods

During the 2016-2019 field seasons, the Paleo-Primate Project Gorongosa discovered and documented seven paleontological localities with fossil vertebrates: GPL-1, GPL-2, GPL-6, GPL-7, GPL-8, GPL-11, and GPL-12. Three additional localities have produced invertebrates only (GPL-3, GPL-9, and GPL-10), and two yielded *ex-situ* stone tools (GPL-4 and GPL-5). Menguere Hill, with abundant fossil wood, is the westernmost fossiliferous locality and it is not identified by a GPL number (Figure 3). These localities are listed in Table 1. This study provides new data and integrates several lines of evidence from the Mazamba Formation, including 1) sedimentology and depositional environments of the fossil localities, 2) radiometric age determinations based on cosmogenic nuclides, 3) stable isotopes from pedogenic carbonates, 4) paleobotanical remains, and 5) vertebrate paleontology. First, we consider each of these lines of evidence separately, emphasizing the materials, analytical methods, and results from each approach. We then synthesize, integrate, and discuss these approaches and their paleoenvironmental and evolutionary significance (for details of each methodological approach see Supporting Information).

### Sedimentology and stratigraphy of the lower Mazamba Formation

Based on regional stratigraphic relationships, sedimentary facies, facies architecture, and the emerging fossil record, Habermann et al. (2019) interpreted the sedimentary successions of the lower member of the Mazamba Formation exposed in the study region as representing a paleoenvironmental mosaic of estuarine and riverine forest/woodland systems. Estuarine sequences accumulated prior to rifting as compound incised-valley fills on a low-gradient coastal plain following transgression, receiving continental sediment from source terranes west of today’s Urema Graben (Habermann et al., 2019). The lower Mazamba successions at the southwestern paleontological sites (GPL-1, GPL-6, GPL-7, GPL-8, GPL-12, see Figure 3) are dominated by basal conglomeratic and sandy facies overlain by clayey sandstones to wackes and sandy clay and marlstone units (Figure 4). These successions are interpreted as lowstand (fluvial) and transgressive (estuarine) assemblages, comprising alluvial channel, bay head delta, shallow central basin or swamp and fluvio-deltaic distributary channel facies from base to top. In contrast, the northeastern localities represent laterally correlative (GPL-9) as well as younger stratigraphic levels (GPL-2, GPL-3); they are sand-dominated and contain marine invertebrates and some fossil mammals. These successions are interpreted as transgressive highstand assemblages consisting of barrier, shore-face, and lagoonal shelf facies (Habermann et al., 2019).

**Figure 4.**
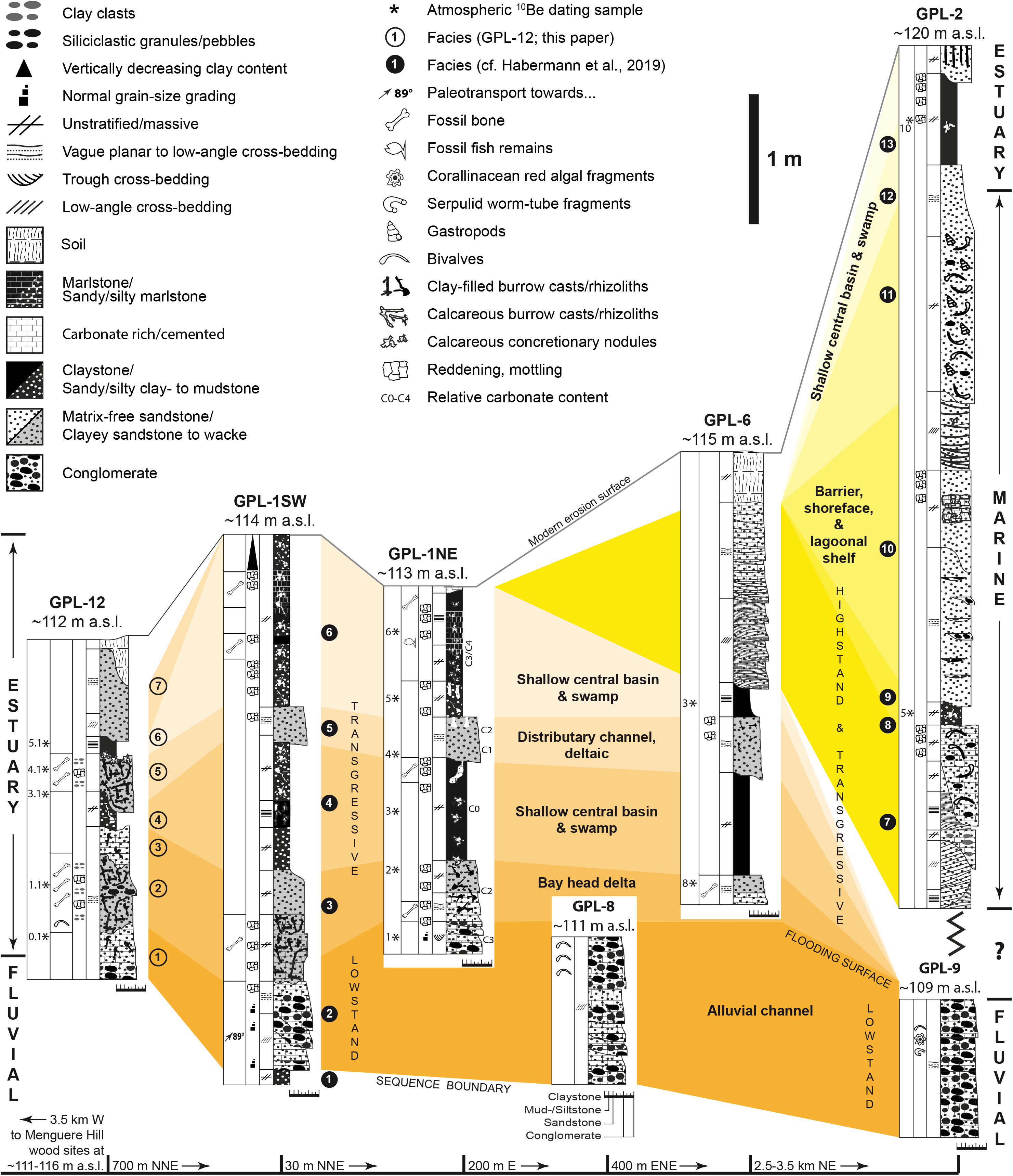
Stratigraphic sections; modified and updated from Habermann et al 2019.

GPL-1 and GPL-12 are the most fossiliferous localities (Supporting Information Figure S1). The sedimentary sequence of GPL-1 was described in detail by Habermann et al. (2019), and here we describe the sedimentary succession of GPL-12 (Figures 3 and 4). The gully sidewall at GPL-12 exposes a 3 m thick section comprising seven distinct sedimentary facies. Coarse, granule- and pebble-bearing quartz sandstones that are moderately cemented by carbonate and contain variable amounts of clay, clayclasts, mottling, and bioturbation form the base of the succession (Facies 1–3). Bedding, occasionally picked out by pebble stringers or abrupt vertical grain-size changes, is only poorly developed. A single cast of a fossil bivalve was found in Facies 2 close to the bottom of the section. Mottling, reddish discoloration, and clay-filled bioturbation casts, including *Thalassinoides* isp., are most common in Facies 2. This facies also yielded numerous vertebrate fossils, which, besides isolated teeth and bone fragments, includes mandibles from various taxa. Brown, sandy claystones with sand-filled bioturbation casts (Facies 4) follow above, which in turn are overlain by clayey sandstones of Facies 5 that represents the second level in the section with large fossil vertebrate remains. At and near the top surface of Facies 5, carbonate accumulated in the form of finely distributed powder, as small concretionary nodules, or as thin crusts, suggesting a disconformity surface. A thin band of olive-green to reddish waxy claystone follows next (Facies 6), which is overlain by medium-grained, well-sorted sandstones that are cross-bedded in places. In Figure 4 we present tentative correlations between localities based on lithological and sedimentological criteria.

Grain-size and sorting characteristics of the basal sandstones of Facies 1–3 suggest a fluvial depositional environment. The vertebrate, invertebrate, and trace fossils in this part of the succession, however, comprise terrestrial and potentially brackish or marine elements (bivalve in Facies 2 as well as *Thalassinoides* isp., most commonly produced by burrowing decapod crustaceans). The fossil remains thus refine paleoenvironmental inferences, suggesting fluvio-deltaic conditions, possibly in a river-dominated estuarine context (bay-head delta assemblage). Fossil preservation and abundance in Facies 2 may suggest high sedimentation rates and relatively rapid burial, perhaps during a storm or flood event. Claystone units in the GPL-12 succession may indicate overbank or mudpond deposition in a fluvio-deltaic environment or may reflect a deepening trend so that estuarine muds formed under brackish to marginal marine conditions following transgression.

### Cosmogenic nuclide dating

To establish a chronology for the Mazamba Formation we applied the authigenic ^10^Be/^9^Be cosmogenic nuclide dating method, hereafter referred to as atmospheric ^10^Be dating, since the method is based on the atmospherically produced isotope ^10^Be (Lebatard et al., 2010). We extracted 15 rock samples from continuous sections measured in the lower member of the Mazamba Formation at GPL-1, GPL-2, GPL-6, and GPL-12 (Figures 3 and 4). To obtain as unaltered and unweathered rocks as possible, samples were taken from freshly excavated trench or section walls. The most fossiliferous and best studied outcrops thus far, GPL-1 and GPL-12, are covered by six and five samples, respectively, that were collected from consecutively younger units present in each section. All sampling positions were documented by total station measurements. Supplementary Table S1 lists all samples collected for dating together with their paleoenvironmental context interpreted from the sedimentary record.

Besides sampling the sedimentary strata to be dated (“fossil samples”), atmospheric ^10^Be dating requires sampling of sediments from modern environments (“modern samples”) equivalent to those reconstructed from the sedimentary record to determine the initial authigenic ratio N_0_ characteristic of the Gorongosa region (Lebatard et al., 2010; Lebatard et al., 2008). To obtain these modern sediment samples, of which we analyzed nine in this study (Supplementary Table S1), a range of environments was sampled, including the banks of three rivers descending from Mount Gorongosa (proximal fluvial settings), the banks of the Pungue and Urema Rivers and the shore of Lake Urema (medial fluvial and lacustrine settings), as well as several localities on the coast, including the Savane River estuary and another estuary northeast of Beira, the shores of which support extensive mangrove swamps and forests (distal coastal, estuarine, and mangrove forest settings). See Supporting Information: Cosmogenic Nuclide Dating for further details.

#### Atmospheric ^10^Be/^9^Be dating – results

The authigenic ^10^Be/^9^Be ratios measured for the modern sediment samples (ranging from 70.9 to 281 x 10^-13^, Supplementary Table S2) are low compared to the range of authigenic ^10^Be/^9^Be ratios of recent surficial continental sediments in general (Graham, Ditchburn, & Whitehead, 2001; Šujan et al., 2016; Wittmann et al., 2012). Due to the dispersion of the obtained N_0_ values (Supplementary Table S2), with a low statistical correlation value, the modern samples were grouped by depositional environments. Then, three scenarios were considered: (1) a direct modern sedimentary/environmental conditions equivalent, (2) a fully estuarine environmental equivalent, and (3) a sedimentary source equivalent. For the first computing (Table 2 part (1)), assuming that the lower Mazamba sediments were deposited in two main paleoenvironments, i.e., fluvio-deltaic and estuarine-lagoonal, we chose modern samples derived from an environmentally equivalent context. For the fossil fluvio-deltaic deposit samples (n=5) (Be18-Gor-GPL1NE-1, Be18-Gor-GPL1NE-2, Be18-Gor-GPL12-0.1, Be18-Gor-GPL12-1.1, and Be18-Gor-GPL12-4.1), data from the modern sample Be18-Bei-EstRi1-1 was used as the N_0_ reference value to calculate depositional ages of 8.6 ± 0.2 Ma and 14.6 ± 0.3 Ma for the first two samples from the base of GPL-1NE. For samples from the basal and middle sections at GPL-12 (GPL12-0.1, -1.1, and -4.1), deposition ages of 17.1 ± 0.5 Ma, 19.5 ± 0.8 Ma, and 16.9 ± 0.6 Ma were calculated, respectively. By contrast, the modern estuarine context samples Be18-Bei-SavEst-1 and Be18-Bei-SavFor-1, for which a weighted mean ^10^Be/^9^Be ratio of 0.640 ± 0.034 x 10^-8^ was obtained, were used as N_0_ reference material to calculate deposition ages for the remaining fossil samples (n = 10) that reflect estuarine-lagoonal conditions. Calculated ages for these samples, coming from middle to upper parts of the GPL-1 and GPL-12 sections, range between 6.9 ± 0.2 Ma (Be18-Gor-GPL1NE-6) and 17.8 ± 0.7 Ma (Be18-Gor-GPL1NE-5).

**Table 2.**
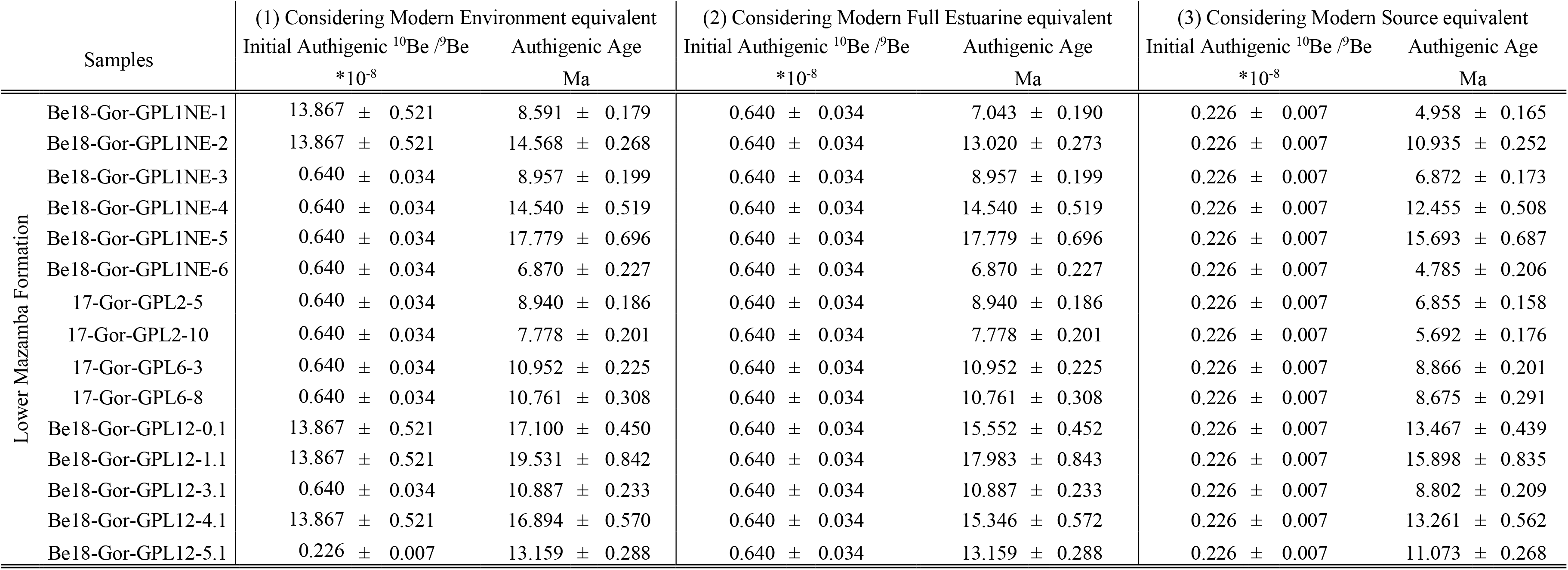
Computed authigenic ages for the lower member of the Mazamba Formation. (1) Modern environmental equivalent sample used for fossil samples Be18-Gor-GPL1NE-1, Be18-Gor-GPL1NE-2, Be18-Gor-GPL12-0.1, Be18-Gor-GPL12-1.1, and Be18-Gor-GPL12-4.1: Be18-Bei-EstRi1-1; Modern environmental equivalent samples used for the other fossil samples: Be18-Bei-SavEst-1 and Be18-Bei-SavFor-1 with a weighted mean ^10^Be/^9^Be ratio of 0.640 ± 0.034 x 10^-8^. (2) Modern estuarine equivalent samples used for all fossil samples: Be18-Bei-SavEst-1 and Be18-Bei-SavFor-1 with a weighted mean ^10^Be/^9^Be ratio of 0.640 ± 0.034 x 10^-8^. (3) Modern source equivalent samples used for all fossil samples: Be18-Gor-Urem-1.1, Be18-Gor-Vun-1.1 and Be18-Gor-VunS1-1.1 with a weighted mean ^10^Be/^9^Be ratio of 0.226 ± 0.007 x 10^-8^.

In the second computing (Table 2 part (2)), assuming the depositional environment for the lower Mazamba Formation was mainly estuarine, only the two modern estuarine context samples (Be18-Bei-SavEst-1 and Be18-Bei-SavFor-1) were considered for age calculations with a mean N_0_ value of 0.64 ± 0.03 x 10^-8^. In this scenario, calculated deposition ages range from 6.9 ± 0.2 Ma (Be18-Gor-GPL1NE-6) to 18.0 ± 0.8 Ma (Be18-Gor-GPL12-1.1) and only the resulting dates for the five fossil fluvio-deltaic samples change with respect to the first computing.

In the third computing (Table 2 part (3)), environmental conditions were largely irrelevant for the choice of modern reference samples. Instead, we chose modern samples for obtaining N_0_ values (mainly for the dissolved ^9^Be input sources) based on sampling localities in the vicinity of the source rocks that the sediments are inferred to be primarily derived from (i.e., Gorongosa Suite granite and gabbro exposed at Mount Gorongosa; Habermann et al., 2019). Matching depositional environments of modern and fossil samples (in this case fluvial) were considered secondarily only in the selection process. The ^10^Be/^9^Be ratios obtained from three modern samples, one from the banks of the Urema River (Be18-Gor-Urem-1.1) and two from the banks of the Vunduzi River (Be18-Gor-Vun-1.1 and Be18-Gor-VunS1-1.1), were used to calculate a weighted mean N_0_ value of 0.226 ± 0.007 x 10^-8^. This weighted mean value was then applied in age calculations to the lower Mazamba samples to be dated. In this approach, resulting ages prove to be slightly younger, ranging between 4.8 ± 0.2 Ma (Be18-Gor-GPL1NE-6) and 15.9 ± 0.8 Ma (Be18-Gor-GPL12-1.1). Thus, under the three different models, all but two of the samples yield dates within the time frame of the Miocene. The lower sections of GPL-12 yield the oldest dates and indicate that the sediments are of early Miocene age. The four samples from GPL-2 and GPL-6 provide late Miocene ages under the three different models. Further information is provided in Supporting Information.

#### Cosmogenic nuclide ^26^Al/^10^Be dating

The upper member of the Mazamba Formation has not yielded any fossils yet, and previous geological work indicates it is much younger than the lower member, but no radiometric dates have been previously reported. We applied the ^26^Al/^10^Be burial dating method based on the decay of ^26^Al and ^10^Be cosmogenic nuclides produced in situ in quartz (SiO_2_) minerals (Lebatard et al., 2014; Lebatard, Bourlès, & Braucher, 2019; Pappu et al., 2011) to date samples from the upper member and thus provide chronological constraints on the fossiliferous lower member. In general, this technique is applicable for the time frame from 100 ka to ∼6 Ma (Granger & Muzikar, 2001). We chose two rock samples collected from two detailed stratigraphic sections in the Mussapassua area in the southeastern corner of GNP where the upper member is well exposed. Under two different models, the samples yielded burial duration dates of ∼1 Ma and indicate that at least part of the upper member is of early Pleistocene age. For analytical details, please see the Supporting Information: Cosmogenic Nuclide Dating as well as Supplementary Tables S3 and S4.

### Pedogenic carbonates

Stable carbon (δ^13^C) and oxygen (δ^18^O) isotope values of 17 pedogenic carbonates from GPL-1 were used to infer regional paleovegetation and climate patterns during the formation of the fossil bearing sediments. δ^13^C values serve as a robust and well-established tool to reconstruct past vegetation growing on the site following soil development. On a continental scale, different biomes differ in the δ^13^C values of associated biomass. Dicots (trees, bushes, herbs) are primarily C_3_ plants, whereas tropical grasses and sedges use the C_4_ photosynthetic pathway and differ in their discrimination against ^13^CO_2_ (Pearcy & Ehleringer, 1984). C_4_ photosynthesis is typically prevalent in warm and seasonally dry, open conditions with high light intensity, whereas the C_3_ pathway is advantageous under low water stress and at high-pCO_2_ conditions. Due to a difference in their discrimination against ^13^C during photosynthesis, δ^13^C values of most C_4_ plants range from -9 to -19 ‰, while those of C_3_ plants lie between -25 and -29 ‰, resulting in ^13^C/^12^C ratios of tropical grasses and sedges ca. 14 ‰ higher than most trees, shrubs, bushes, and herbaceous plants (Cerling, Harris, Ambrose, Leakey, & Solounias, 1997). The comparably small variability of δ^13^C in C_4_ plants can be attributed to three different C_4_ photosynthetic subpathways (Pearcy & Ehleringer, 1984), while the variation in δ^13^C among C_3_ plants is affected by a variety of environmental factors including trophic effect, precipitation, temperature, drought, canopy density, salinity, light intensity, nutrient levels, and partial pressure of CO_2_ (Diefendorf, Mueller, Wing, Koch, & Freeman, 2010; Ehleringer & Monson, 1993; Farquhar, Ehleringer, & Hubick, 1989; Kohn, 2010; E. Medina & Minchin, 1980; Ernesto Medina, Montes, Guevas, & Rokzandic, 1986). Collectively, however, these effects on δ^13^C of C_3_ plants are still considerably small compared to the differences between C_3_ and C_4_ biomass. Pedogenic carbonate formed in equilibrium with soil-respired CO_2_ is typically enriched in ^13^C by 13.5 to 17.0 ‰ compared to the CO_2_ which respired from plants or was released during decomposition of soil organic carbon and related organic matter (Cerling, 1984; Cerling, Wang, & Quade, 1993).

Pedogenic carbonate forms in oxygen isotope equilibrium with soil water (Cerling & Quade, 1993). The δ^18^O value of soil carbonate is a function of soil water composition and temperature. Soil water is derived from meteoric water, but can differ from this source water due to enrichment through evaporation from the soil surface, mixing with (evaporatively ^18^O-enriched) infiltrating water, and/or the addition of isotopically distinct water from overland and vadose zone flow (Hsieh, Chadwick, Kelly, & Savin, 1998). Nevertheless, δ^18^O values of modern pedogenic carbonate have a strong positive correlation with the composition of meteoric water, which in turn has a positive correlation with local air temperature (Rozanski, Araguás-Araguás, & Gonfiantini, 1992). Collectively, this makes paleosol carbonate an important paleoclimate proxy. The composition of local meteoric water has a large influence on δ^18^O of soil water and hence pedogenic carbonate δ^18^O. Today, the climate of central Mozambique is a result of interactions between the African Monsoon, the Intertropical Convergence Zone, and the Zaire Air Boundary. These complex patterns complicate the comparison of absolute δ^18^O values of distant localities, due to possibly different isotopic composition of local precipitation. See Supporting Information: Pedogenic Carbonates for methodological details.

#### Pedogenic carbonates – results

All results are listed in Table 3 and shown in Figure 5. Stable carbon isotope ratios of pedogenic carbonates of GPL-1 vary between -9.3 and -5.9 ‰ with an average value of -7.3 ± 1.0 ‰, while oxygen isotopes ratios fluctuate from 25.4 to 26.5 ‰ with an average of 25.9 ± 0.3 ‰. There is very low correlation between δ^13^C and δ^18^O present (R^2^ = 0.1). Overall stratigraphic trends cannot be detected in either of the two datasets. Carbonate content of the nodules is generally >50 % with only one sample having a significantly lower carbonate content (16 %), but comparable isotopic values. The average carbonate content is 80 ± 20 %.

**Table 3.** Stable carbon and oxygen isotopic values with sample ID, distance from the base of section GPL-1NE, amount of untreated carbonate powder and carbonate content. For stratigraphic context see Habermann et al 2019 and Figure 4.

| Sample ID | Distance from<br>base [cm] | $\delta^{13}\text{C}_{\text{VPDB}}$<br>[‰] | $\delta^{18}\text{O}_{\text{VSMOW}}$<br>[‰] | Weight<br>[μg] | Carbonate<br>content [%] |
| --- | --- | --- | --- | --- | --- |
| GLP1-1NE-25 | 390 | -6.8 | 26.2 | 112 | 91 |
| GLP1-1NE-24 | 385 | -9.1 | 25.8 | 144 | 87 |
| GLP1-1NE-23 | 380 | -7.7 | 26.4 | 121 | 89 |
| GLP1-1NE-22 | 360 | -8.6 | 26.0 | 129 | 95 |
| GPL1-1NE-21 | 350 | -7.1 | 25.7 | 366 | 16 |
| GLP1-1NE-19 | 340 | -6.4 | 25.7 | 146 | 96 |
| GLP1-1NE-18 | 335 | -7.0 | 26.0 | 135 | 91 |
| GLP1-1NE-17 | 330 | -7.0 | 25.6 | 135 | 93 |
| GLP1-1NE-16 | 325 | -7.5 | 25.5 | 170 | 89 |
| GLP1-1NE-15 | 320 | -6.7 | 26.1 | 139 | 88 |
| GLP1-1NE-14 | 310 | -9.3 | 25.9 | 159 | 78 |
| GPL1-1NE-13 | 200 | -7.6 | 25.7 | 149 | 83 |
| GPL1-1NE-10 | 145 | -7.4 | 25.4 | 123 | 50 |
| GPL1-1NE-09 | 135 | -7.5 | 26.0 | 131 | 62 |
| GPL1-1NE-08 | 120 | -6.3 | 26.2 | 136 | 79 |
| GPL1-1NE-07 | 110 | -5.9 | 26.5 | 131 | 86 |
| GPL1-1NE-06 | 90 | -6.3 | 26.3 | 150 | 80 |

**Figure 5.**
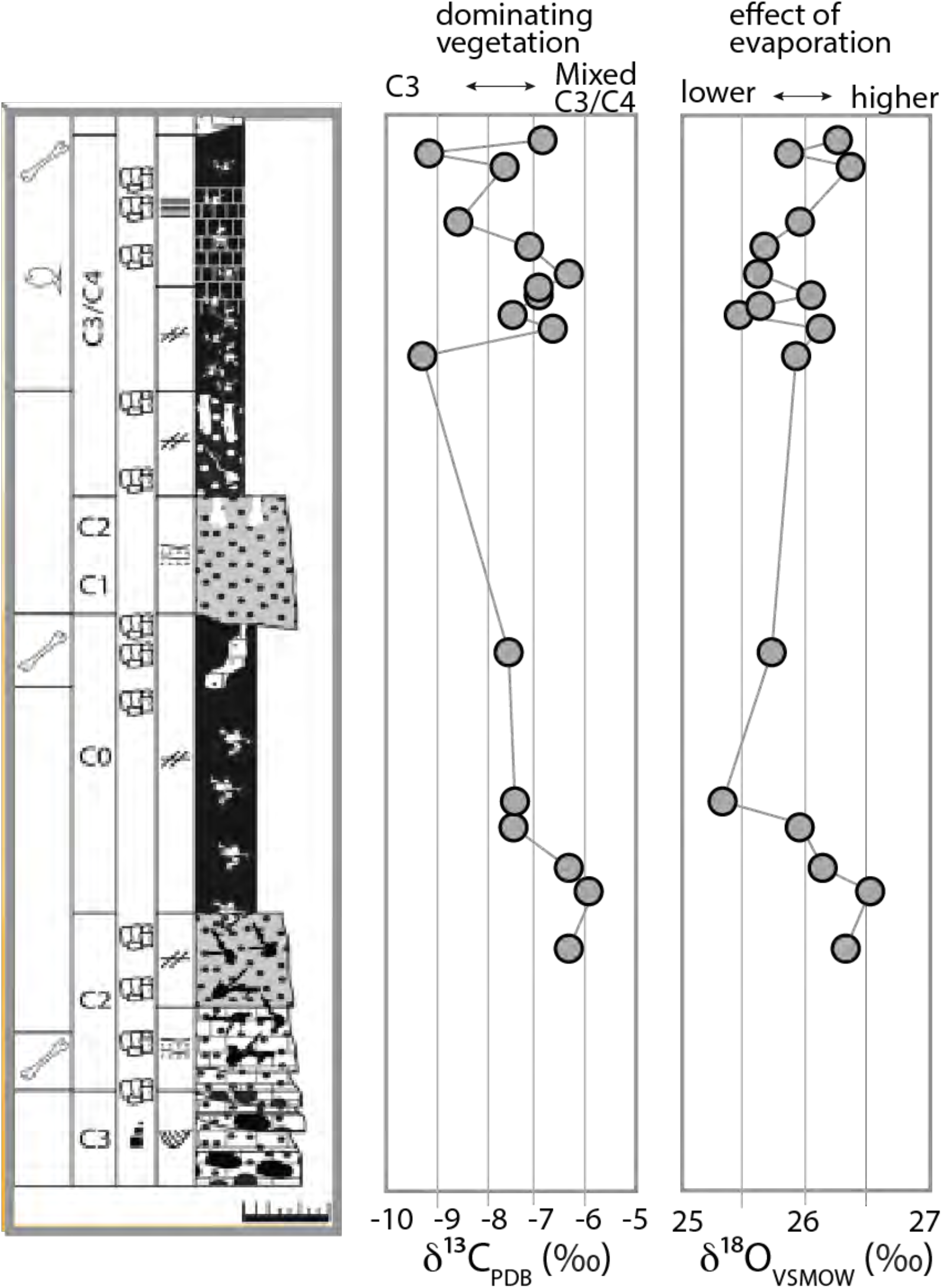
Stable carbon, δ^13^C, and oxygen, δ^18^O, isotopic compositions related to the stratigraphic column of GPL-1NE (see Habermann et al., 2019).

Carbon isotope values average -7.3 ± 1.0 ‰ and never exceed -5.9 ‰. Such low values are typical for C_3_ dominated ecosystems characterized by woodland, bushland, or wooded grassland environment with a mix of C_3_/C_4_ vegetation. Following the vegetation classification of (White, 1983), this would indicate average woody cover of at least 50 % (for the average δ^13^C value of -7.3 ‰), using the ‘paleo-shade’ proxy (Cerling et al., 2011). The oxygen isotopic values of pedogenic carbonates from the paleosols of GPL-1 show fluctuations of only 1.1 ‰ towards a relatively persistent climate with no large variation in temperature, source water supply or effects of evaporation. Without constraints on paleotemperature or ancient soil water oxygen isotopic composition, temporal and geographic variations in fossil soil carbonate δ^18^O values can only be used to identify qualitative changes in climatic patterns, but the relatively low δ^18^O values could indicate a mesic climate with high water supply, which is also supported by the sedimentology, geology and fossil faunal and floral assemblages of this costal riverine forest/woodland ecosystem (Habermann et al., 2019).

### Paleobotany

At Menguere Hill, about 3.5 km west of GPL-1, there are large, silicified tree trunks (Figure 6) measuring up to 1.6 m in diameter, as well as scattered fragments of fossil wood, as noted by Pickford (2013) and described by Habermann et al. (2019). Menguere Hill rises 40 m above the surrounding landscapes and exposes a series of silicified limestone beds. During the 2016-2018 field seasons, we collected 41 specimens of well-preserved fossil wood for microscopic analysis of thin sections and here we present a preliminary taxonomic list and the paleoecological implications of the taxa. The Gorongosa sample includes the palm *Hyphaene* (Palmae, family Arecaceae), which is widespread in the humid, hot lowlands of tropical Africa. The most abundant taxon in the collection is *Entandrophragmoxylon* (African mahogany, family Meliaceae) (Figure 7). The modern genus *Entandrophragma* is restricted to tropical Africa, and some species can reach up to 60 m in height. We have previously reported the presence of *Terminalioxylon* (family Combretaceae) (Habermann et al., 2019), a genus that is most diverse in bushveld and savannas, and includes some mangrove species. There are also samples of *Zizyphus* (family Rhamnaceae), which is common along watercourses, and *Zanha* (family Sapindaceae), found in open woodland to dense ravines and riverine forests (Arbonnier, 2004; Beentje, 1994; Coates Palgrave, 2002). A further observation to note is that cross sections of the wood vessels indicate mesophytic trees that cannot tolerate water stress. We interpret the Menguere Hill succession as a correlative inland equivalent to the estuarine fossil sites farther to the east based on similar elevations (see Table 1) (Habermann et al., 2019).

**Figure 6.**
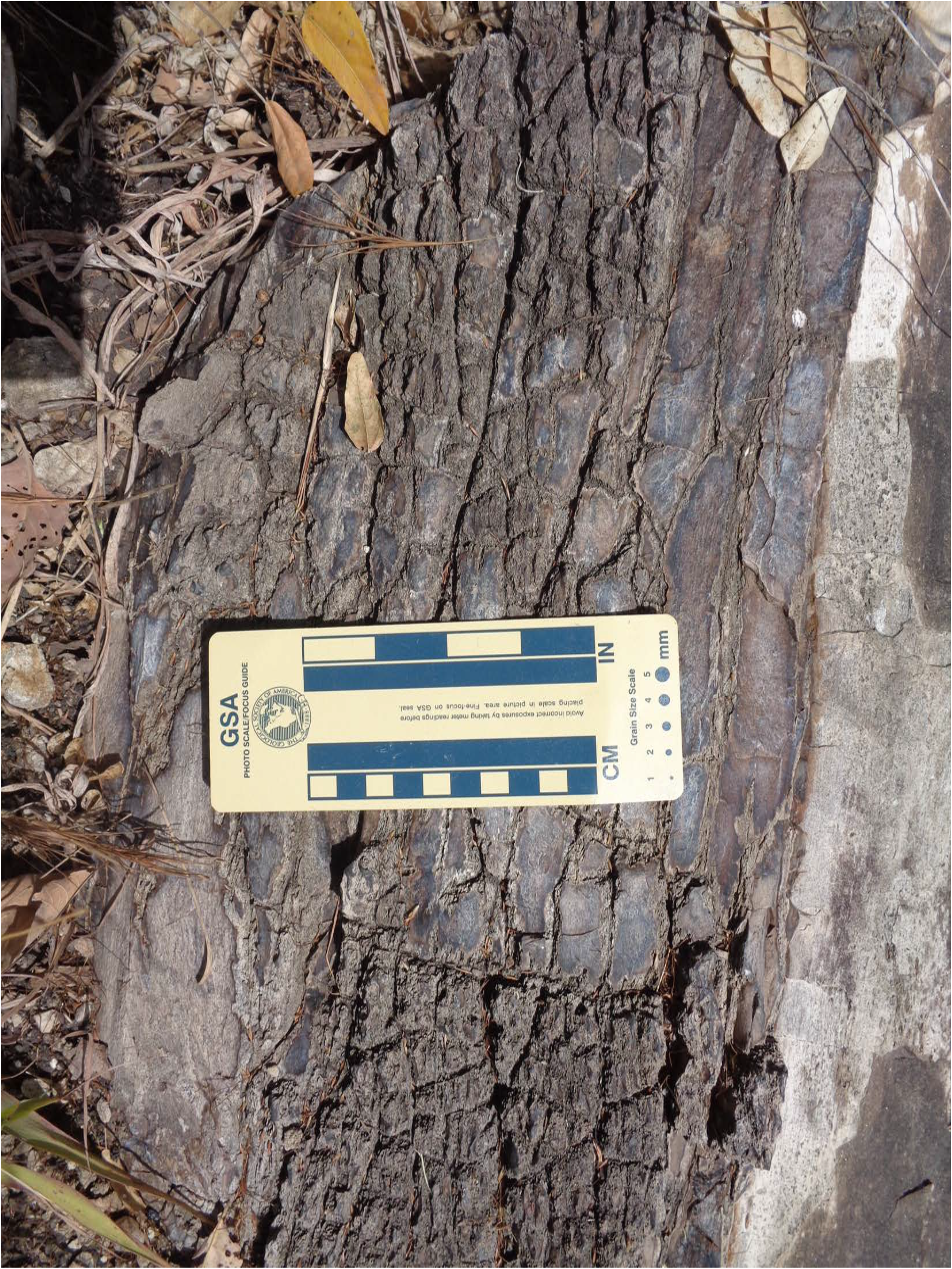
Silicified tree trunk with bark preserved at Menguere Hill.

**Figure 7.**
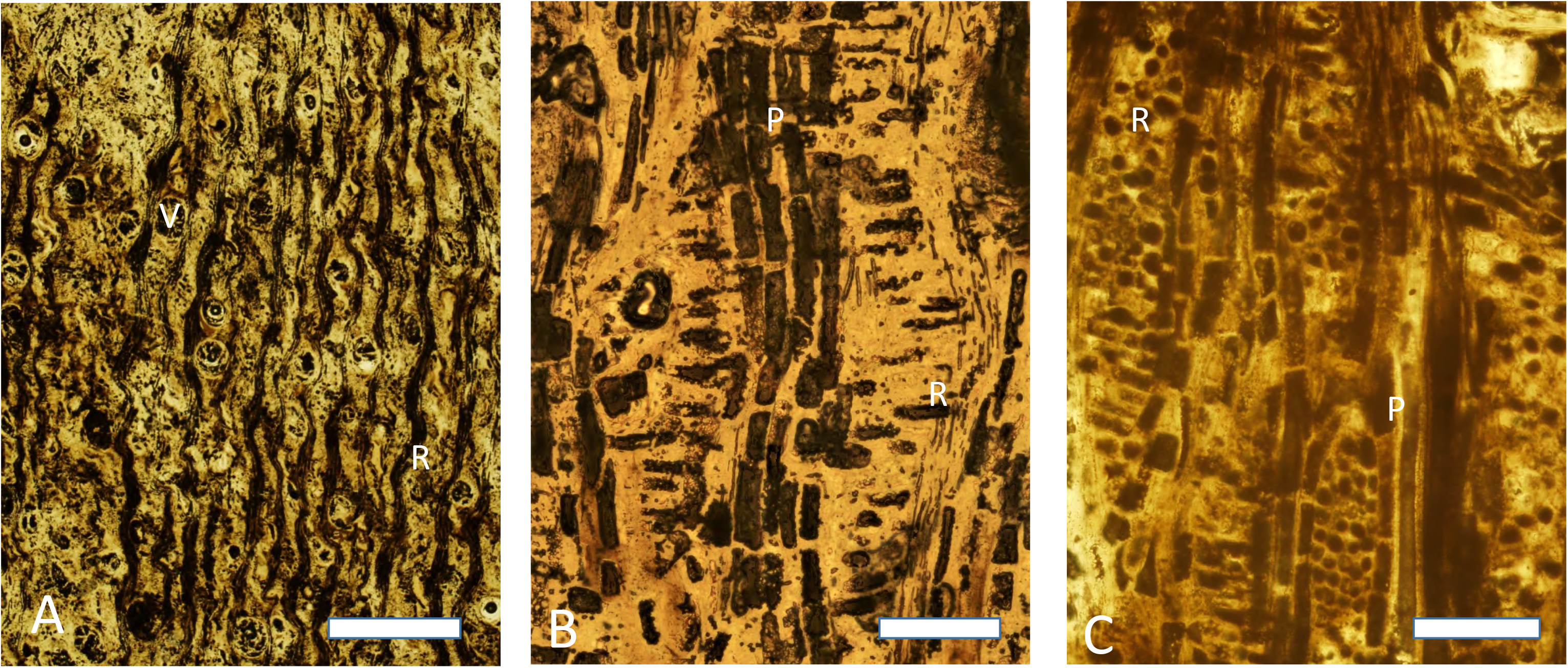
Photomicrographs of thin sections of fossil wood specimen PPP-G-36 from Menguere Hill, *Entandrophragmoxylon* sp. (Meliaceae, African Mahogany). **A)** Transverse section showing large mostly solitary vessels, vasicentric to aliform parenchyma and wide rays with dark contents. **B)** Radial longitudinal section with a vertical column of axial parenchyma cells, and horizontal radial parenchyma cells that are procumbent. **C)** Tangential longitudinal section with vertical columns of axial parenchyma cells and lens-shaped outline of rays with circular parenchyma cells. Letters: V = vessel; R = ray; P = axial parenchyma. Scale bars: A = 1cm; B, C = 500µm.

### Systematic paleontology

Here we describe several key specimens from the lower Mazamba Formation. Additional Gorongosa fossils have recently been excavated and are currently under preparation, curation, and study (see Supporting Information).

Class Chondrichthyes Huxley, 1880
Subclass Elasmobranchii Bonaparte, 1838
Order Carcharhiniformes Compagno, 1977
Family Carcharhinidae Jordan & Evermann, 1896
Genus *Galeocerdo* Müller & Henle, 1837
Referred specimens: PPG2017-P-121 from GPL-1, PPG2018-P-224 from GPL-1, PPG2019-P-126, 129, 176 from GPL-12
*Galeocerdo aduncus* Agassiz, 1843
Referred specimen: PPG2019-P-127 from GPL-12

Six specimens of shark teeth have been recovered from the Gorongosa sedimentary sequence. Four of these are fragmentary teeth from GPL-1 (PPG2017-P-121, PPG2018-P-224) and GPL-12 (PPG2019-P-126, PPG2019-P-127), and two are complete crowns and roots from GPL-12 (PPG2019-P-127, PPG2019-P-129) (Figure 8). For shark teeth we use the terminology of Türtscher et al. 2021. The following descriptions and analyses are based on the two complete teeth. One of these teeth (PPG2019-P-129), however, has some weathering on the apex that removed part of the distal cutting edge. The apex of the Gorongosa teeth is dominated by a primary cusp that leans distally. Serrations are present in the mesial cutting edge and the distal heel, but only lightly developed or absent along the apex. The mesial cutting edge has more than a dozen primary serrations that decrease in size away from the apex. The heel is relatively straight and with ∼10 primary serrations decreasing in size distally. The serrations are simple (not compound), with only primary serrations visible (no secondary serrations). The outline of the mesial cutting edge has a distinct break between the apex and the rest of the serrated mesial cutting edge with two lines meeting at an obtuse angle (140° in PPG2019-P-127 and 155° in PPG2018-P-129). The length of the apex is one third or less of the length of the rest of the mesial cutting edge. The mesiodistal length of the tooth exceeds its height. The root is relatively thick, bilobate and well-arched, with the slightly asymmetrical lobes forming an obtuse angle. The six specimens differ in coloration, weathering, and preservation, and appear to represent distinct individuals deriving from two localities separated by ∼700 m. In overall characteristics, the shark teeth have the cockscomb appearance typical of the genus *Galeocerdo*, tiger sharks.

**Figure 8.**
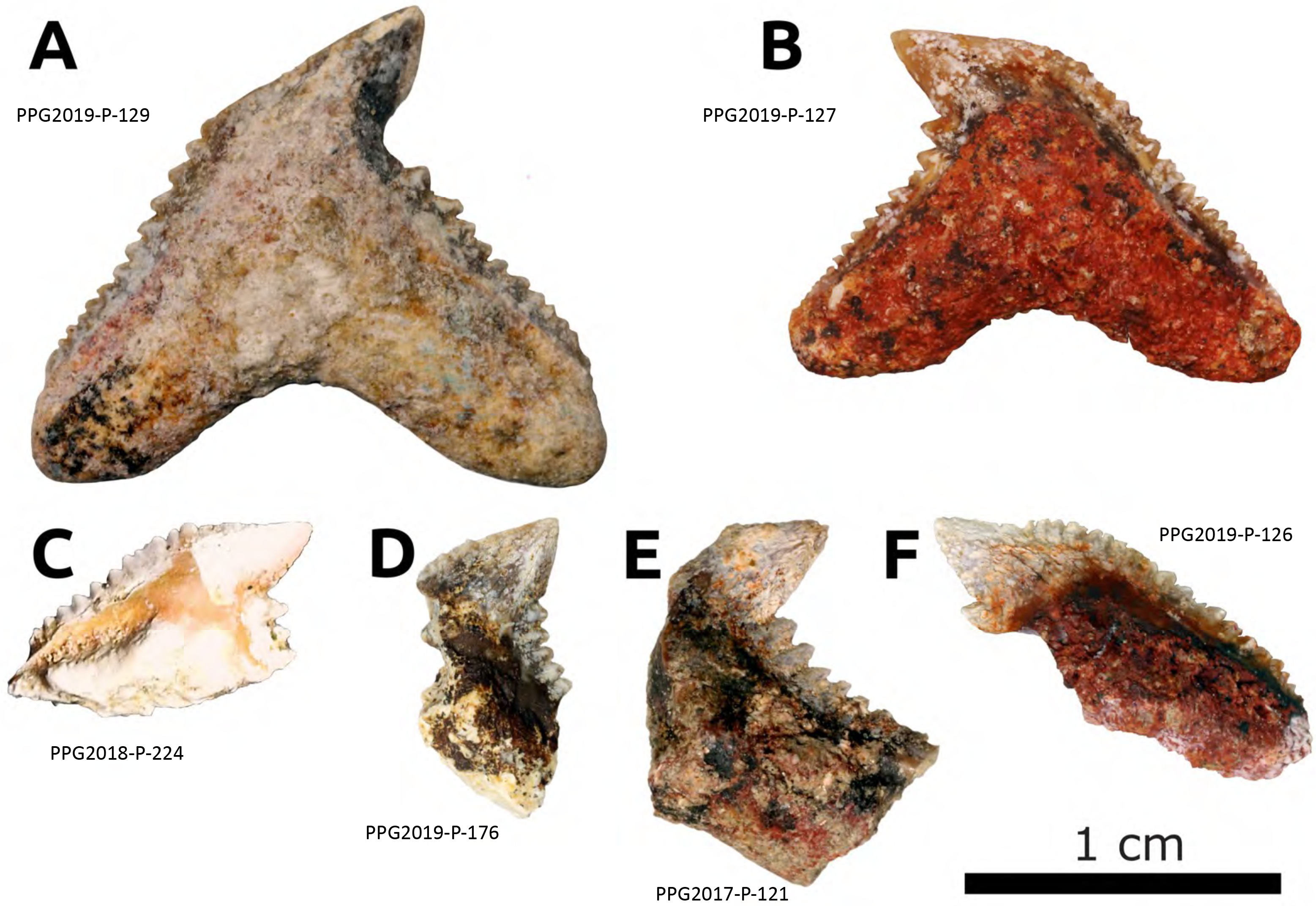
Gorongosa fossil sharks, all in the genus *Galeocerdo*, tiger sharks. **A)** PPG2019-P-129; **B)** PPG2019-P-127; **C)** PPG2018-P-224; D) PPG2019-P-176; E) PPG2017-P-121; F) PPG2019-P-126.

To assess the taxonomic affinities of the Gorongosa shark specimens, we carried out a series of 2D morphometric analyses of the two complete specimens. We compiled a set of fossil shark photographs from the existing literature to obtain a suitable comparative sample of 600 specimens (Supplementary Table S5). From this comparative sample we used three datasets including: 1) all 600 specimens from four different genera (*Galeocerdo*, *Physogaleus*, *Carcharhinus*, and *Hemipristis*), 2) a subset of 547 specimens from species of *Galeocerdo* and *Physogaleus*, 3) a subset including 436 specimens from different species of the genus *Galeocerdo*. We carried out Principal Component Analyses (PCA) of these datasets followed by multi-group Linear Discriminant Analyses (LDA) to classify the Gorongosa specimens into taxonomic categories (see Supporting Information: Morphometric Analysis: Chondrichthyes for methodological details).

The first PCA considering four genera of sharks shows that both Gorongosa specimens are located within the convex hulls of *Galeocerdo* (Figure 9A). In the second PCA, considering eight species of *Galeocerdo* and *Physogaleus*, Gorongosa B is located near the center of the *Galeocerdo aduncus* convex hull, whilst Gorongosa A is in a marginal position near the edges of *G. cuvier* and *G. capellini* (Figure 9B). In the third PCA, which considers only species of *Galeocerdo*, Gorongosa B is again near the center of the *G. aduncus* convex hull, whilst Gorongosa A is near the edges of *G. cuvier* and *G. capellini* (Figure 9C). The three LDA models using the Principal Components (PCs) that accounted for 90% of the variance of the sample clearly distinguishes among the taxonomic categories, displaying good performances with satisfactory classification results after cross-validation (Supplementary Table S6). When using the obtained discriminant functions to classify the Gorongosa fossil sharks into these taxonomic categories (as a way of assessing morphological affinities), they were robustly classified within the genus *Galeocerdo*. When classifying the fossils using the species categories, Gorongosa A was classified within *Galeocerdo cuvier*, whilst Gorongosa B was strongly categorized within *Galeocerdo aduncus*. The species *G. aduncus* has a temporal span from the Oligocene to the end of the Miocene, whilst the earliest record of the extant species *G. cuvier* is from the Middle Miocene (Soto Ovalle, 2016; Türtscher et al., 2021).

**Figure 9.**
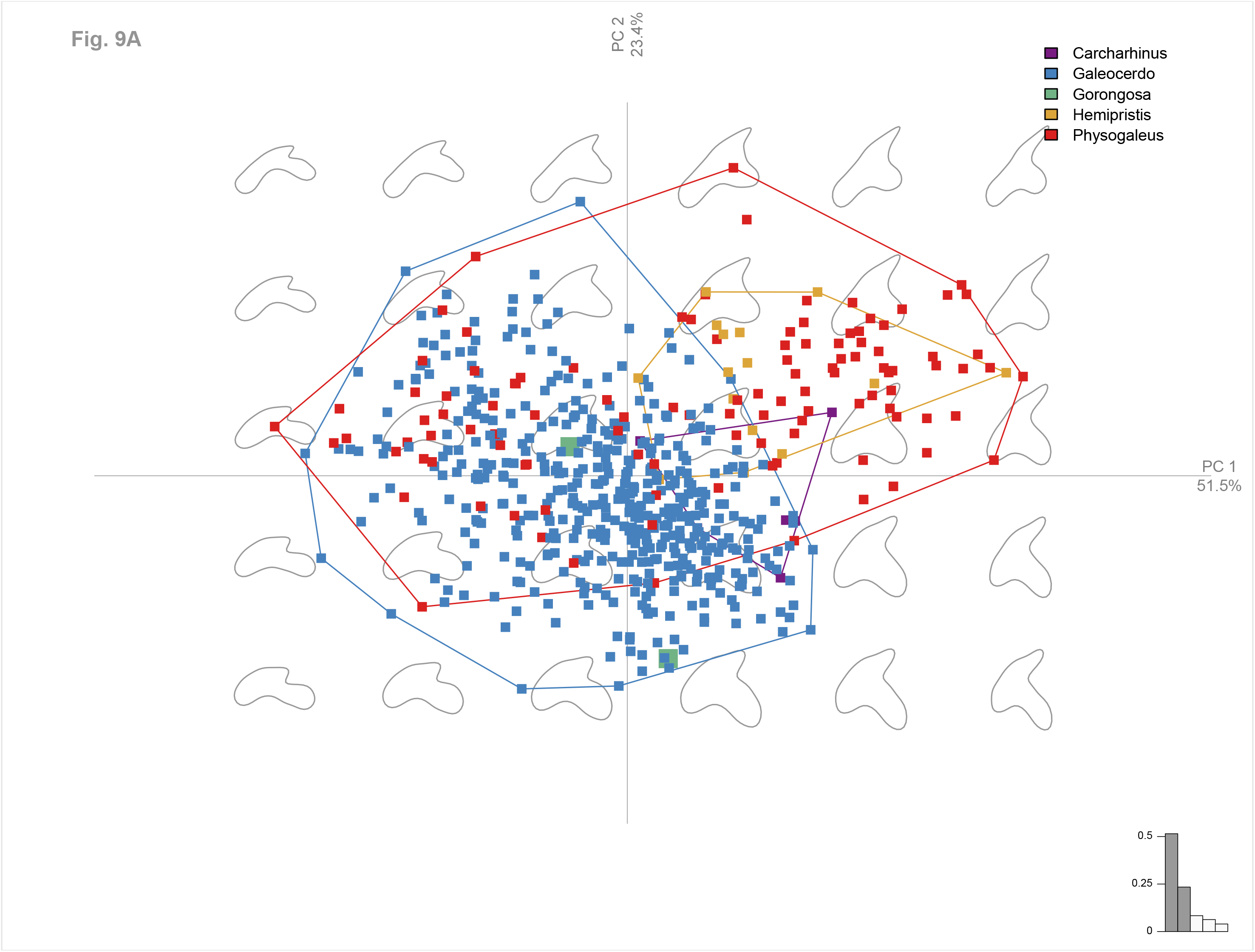
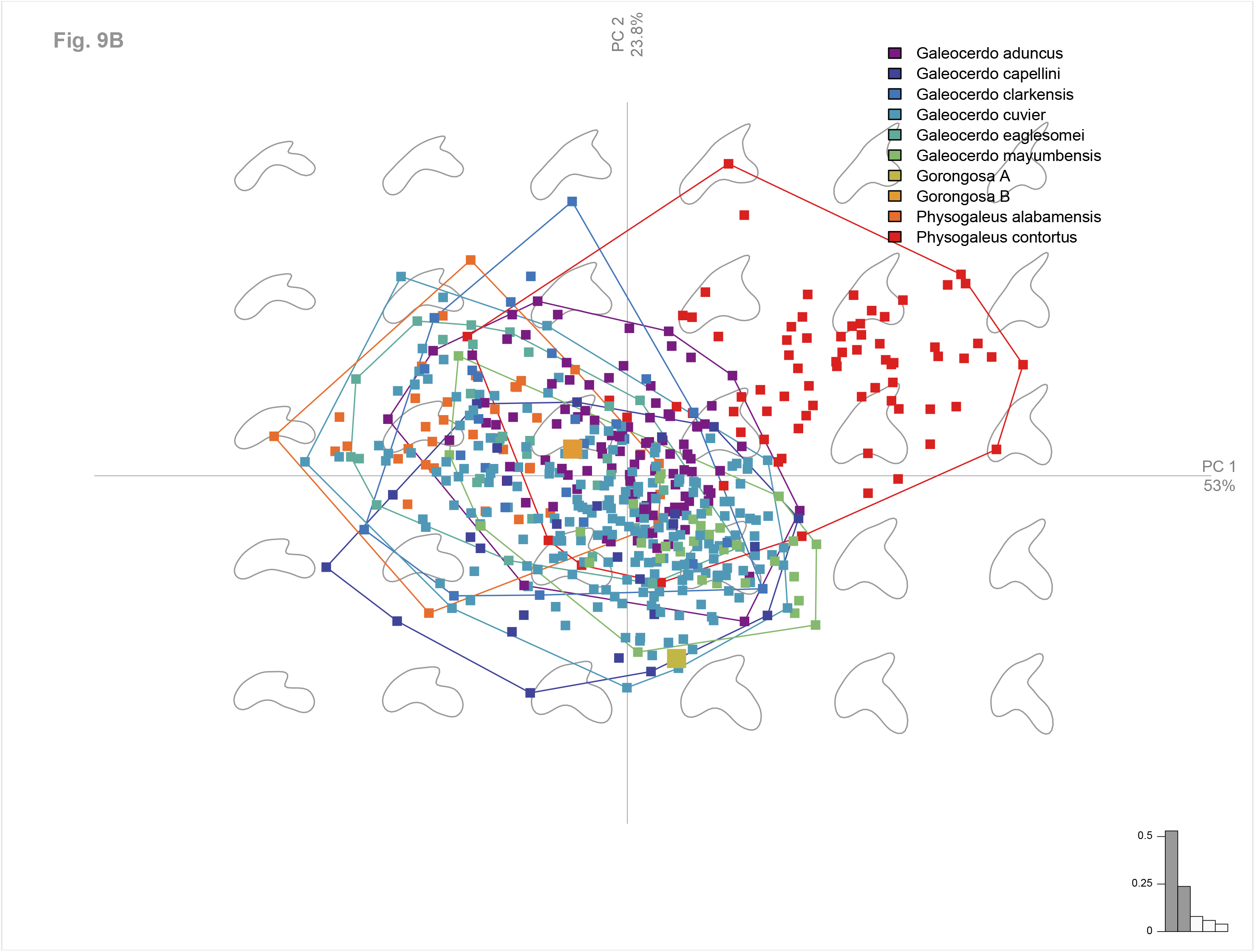
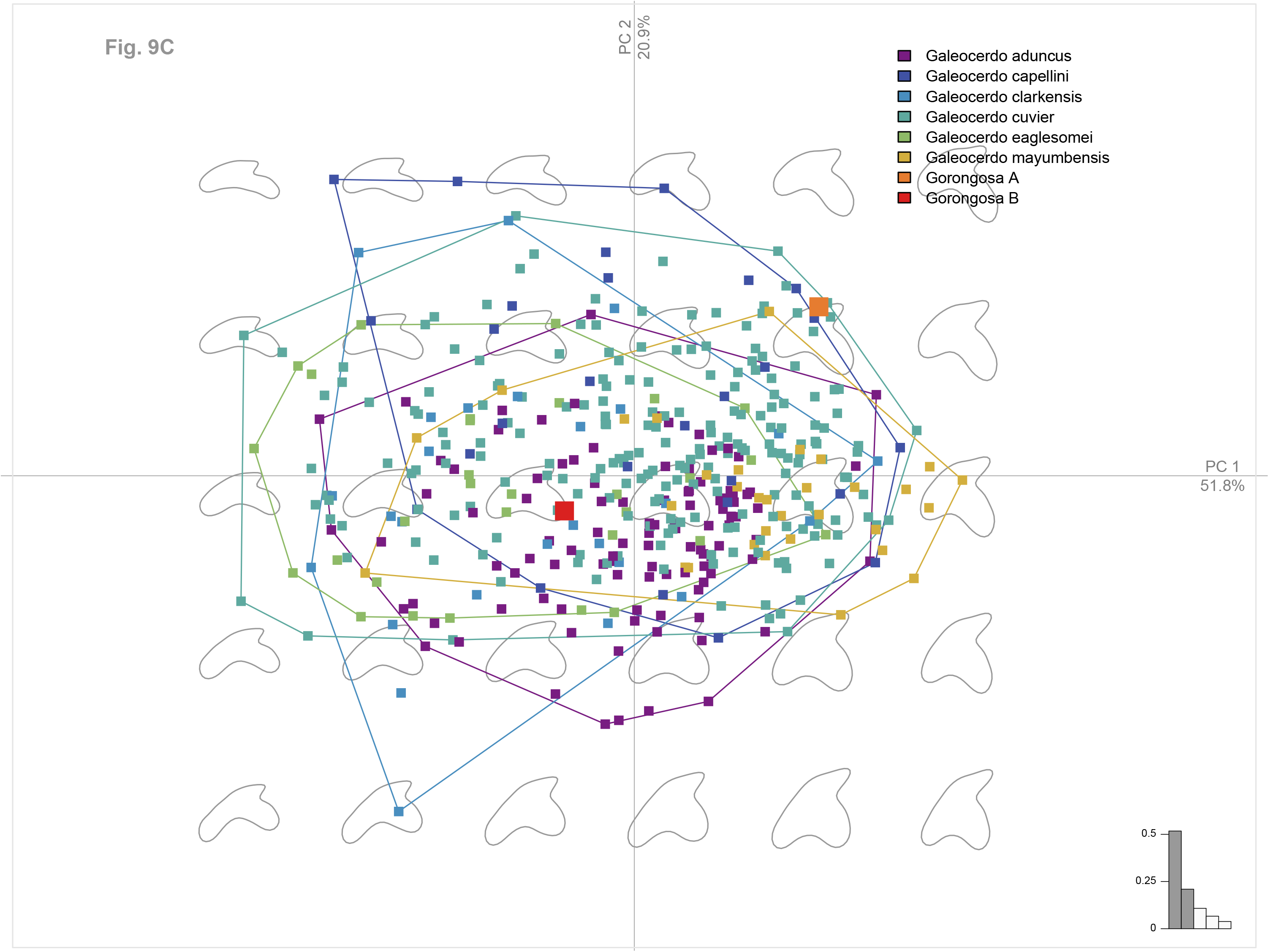
**A)** PCA of 600 Miocene shark teeth from the genera *Carcharhinus*, *Galeocerdo*, *Hemipristis*, and *Physogaleus*, and including the two Gorongosa complete crowns. **B)** PCA of 547 Miocene shark teeth of the species *Galeocerdo* sp., and *Physogaleus* sp., and the Gorongosa specimens. **C)** PCA of shark teeth including the species *G. aduncus, G. capellini, G. clarkensis, G. cuvier, G. eaglesomei,* and *G. mayumbensis*, with the Gorongosa specimens.

The size and morphology of the fragmentary teeth in the Gorongosa collection is consistent with those of the complete crowns, and we attribute all six specimens to the same genus. *Galeocerdo* upper and lower teeth are very similar, but they increase in breadth relative to height posteriorly. The teeth of juvenile tiger sharks have fewer serrations than those of adults (Türtscher et al., 2021). The Gorongosa fossil teeth are functionally similar to those of the extant tiger shark, and we may infer similar function in piercing large prey.

Batoidea Compagno, 1973
Order Myliobatiformes Compagno, 1973
Referred specimen: PPG2018-P-257 from GPL-1

A single fragment of batoid symphyseal teeth was found at GPL-1. This indicates that at least two taxa of cartilaginous fishes occur in the Gorongosa fossil record, one species of shark and one species of ray. Most batoid species live in tropical and subtropical coastal waters, and some can occur in estuaries.

Order Testudines Batsch, 1788
Referred specimens: PPG2016-P-12, 13, 14, 27, 55, PPG2017-P-42, 44, 87, 95, PPG2018-P-10, 201, 203, 206, 217, 233, 234, 235, 270, 271
Family Testudinidae, 1788
Referred specimen: PPG2016-P-9

There are 20 specimens of turtles and tortoises in the Gorongosa fossil collections, which include fragments of carapace and plastron. One of the first specimens to be recovered in the field was PPG2016-P-9, a plastron fragment consistent in thickness and morphology with terrestrial tortoises (family Testudinidae) (Figure 10A), which have been present in Africa since the late Eocene (Holroyd & Parham, 2003; Zouhri et al., 2017). Most specimens are fragmentary but further analyses will aim to refine the taxonomic attributions.

Order Crocodylia Gmelin, 1789
Family Crocodylidae Cuvier, 1807
Crocodylidae indet.
Referred specimens: PPG2016-P-10, 23, PPGG2017-P-43, 49, 73, 80, 89, PPG2018-P-100, 161, 162, 222, 223, 241, 252, 264, PPG2019-P-116, 117, 128

**Figure 10:**
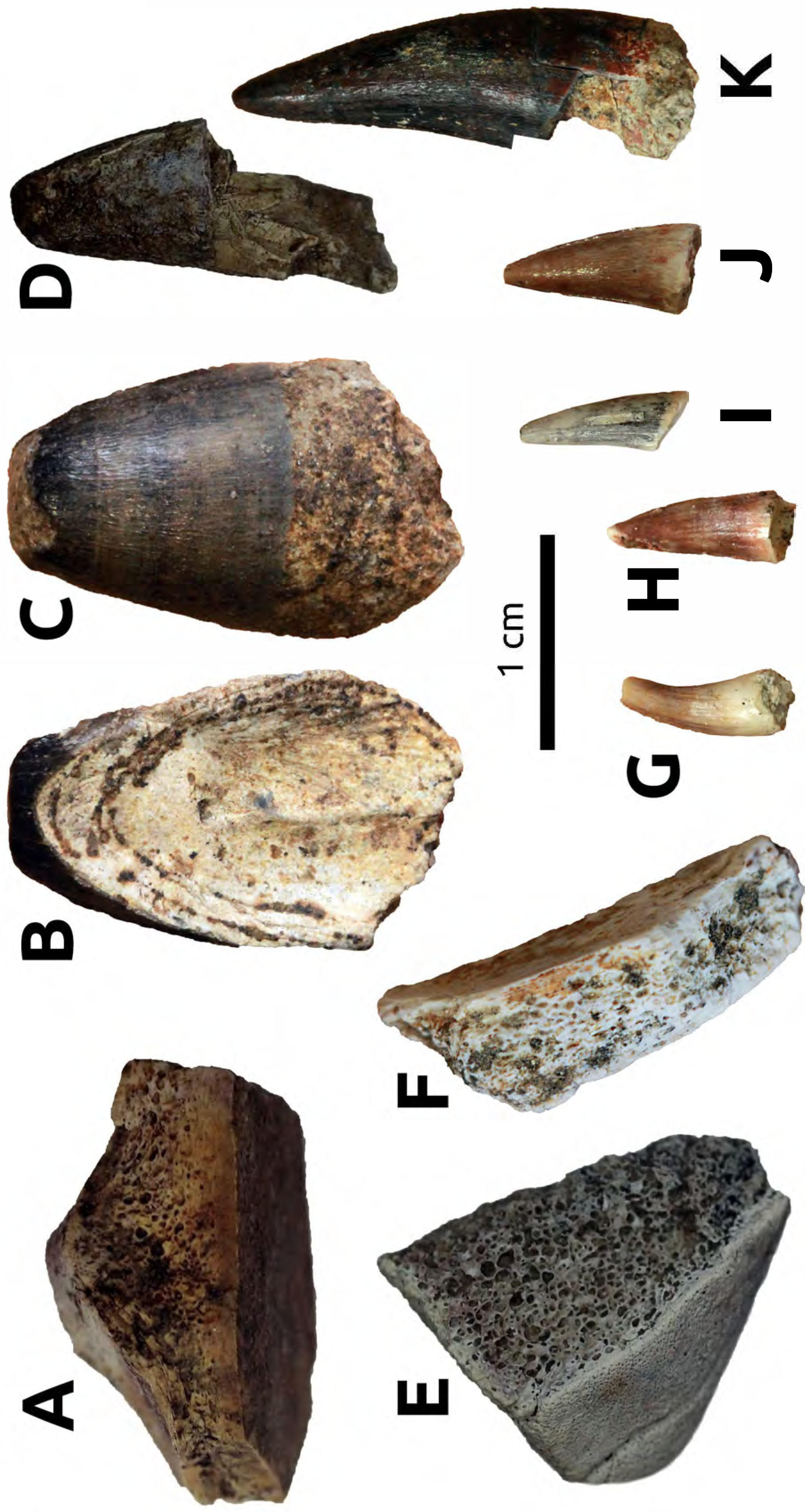
Some fossil reptiles from Gorongosa. **A)** Testudines, **B)** Crocodylia.

There are 18 teeth and tooth fragments attributed to Crocodylidae. Their abundance attests to relatively stable bodies of water in the region. Tooth crown morphologies are consistent with size and shape heterodonty in brevirostrine taxa (Figure 10B) (D’Amore, Harmon, Drumheller, & Testin, 2019; Kieser, Klapsidis, Law, & Marion, 1993). Although represented by small sample sizes, maximum tooth crown lengths indicate body sizes similar to comparatively small-bodied crocodylids from the Paleogene and early to middle Miocene of North African and sub-Saharan formations (Brochu & Gingerich, 2000; Conrad et al., 2013; Cossette et al., 2020), as opposed to the gigantic late Miocene-Pleistocene taxa from East Africa (Brochu & Storrs, 2012; Storrs, 2003). A single broken, poorly preserved tooth is elongate and slightly recurved distally, similar to the condition in longirostrine, piscivorous tomistomine and gavialoid taxa, suggesting the presence of at least two crocodylid taxa in the lower member of the Mazamba Formation.

Mammalia Linnaeus, 1758
Afrotheria Stanhope et al., 1998
Order Hyracoidea Huxley, 1869
Family Saghatheriidae Andrews, 1906
Referred specimens: PPG2018-P-1, 2

Hyraxes (order Hyracoidea) belong to the Afrotheria, a clade of mammals with deep evolutionary roots in Africa. There are five species of modern hyraxes, all in the family Procaviidae, but in the past there were at least four additional families: Geniohyidae, Saghatheriidae, Titanohyracidae, and Pliohyracidae. Hyracoids in the Paleogene of Africa were abundant and diverse, both taxonomically and functionally, but declined in overall diversity during the late Miocene (Rasmussen & Gutiérrez, 2010).

The Gorongosa sample includes an individual with left and right mandibular fragments (Figure 11) excavated in situ from Facies 2 at GPL-12 (Supporting Information Figure S1). The hyracoid mandibles represent some of the oldest mammals found so far in the Gorongosa sequence (early Miocene based on the atmospheric ^10^Be dates). The left hemimandible (PPG2018-P-1) has the complete premolar-molar dentition, from p1 to m3, but the specimen is extremely fragile, so it remains in its plaster jacket for protection and only the buccal and occlusal aspects are visible. The right mandible fragment (PPG2018-P-2) has a complete set of molars (m1-m3) and two detached premolars (p3 and p4). Tooth measurements are given in Table 4. The mandibular body, as seen on the left side, shows a slight depression on the buccal side below the level of m1-m2. The cheek teeth increase monotonically in mesio-distal length from p1 (12.81 mm) to m3 (31.01 mm). The teeth are brachydont, and the molars are bilophodont with well-developed transverse crests. The posterior premolars, p3-p4, are molarized. In the molars, the protoconid is large and gives rise to the protocristid that extends to the metaconid and forms the mesial loph. The paraconid is reduced and the metaconid is the tallest cusp. The hypoconid gives rise to a marked hypocristid that extends to the entoconid and forms the distal loph. The m3 has a well-developed hypoconulid and a third loph joins the hypoconulid with the endoconulid. A continuous cingulum occurs along the mesial, buccal, and distal parts of the molars. The well-developed transverse crests and the low-crowned molars of the Gorongosa specimens most likely indicate a folivorous diet based on soft leaves.

**Figure 11.**
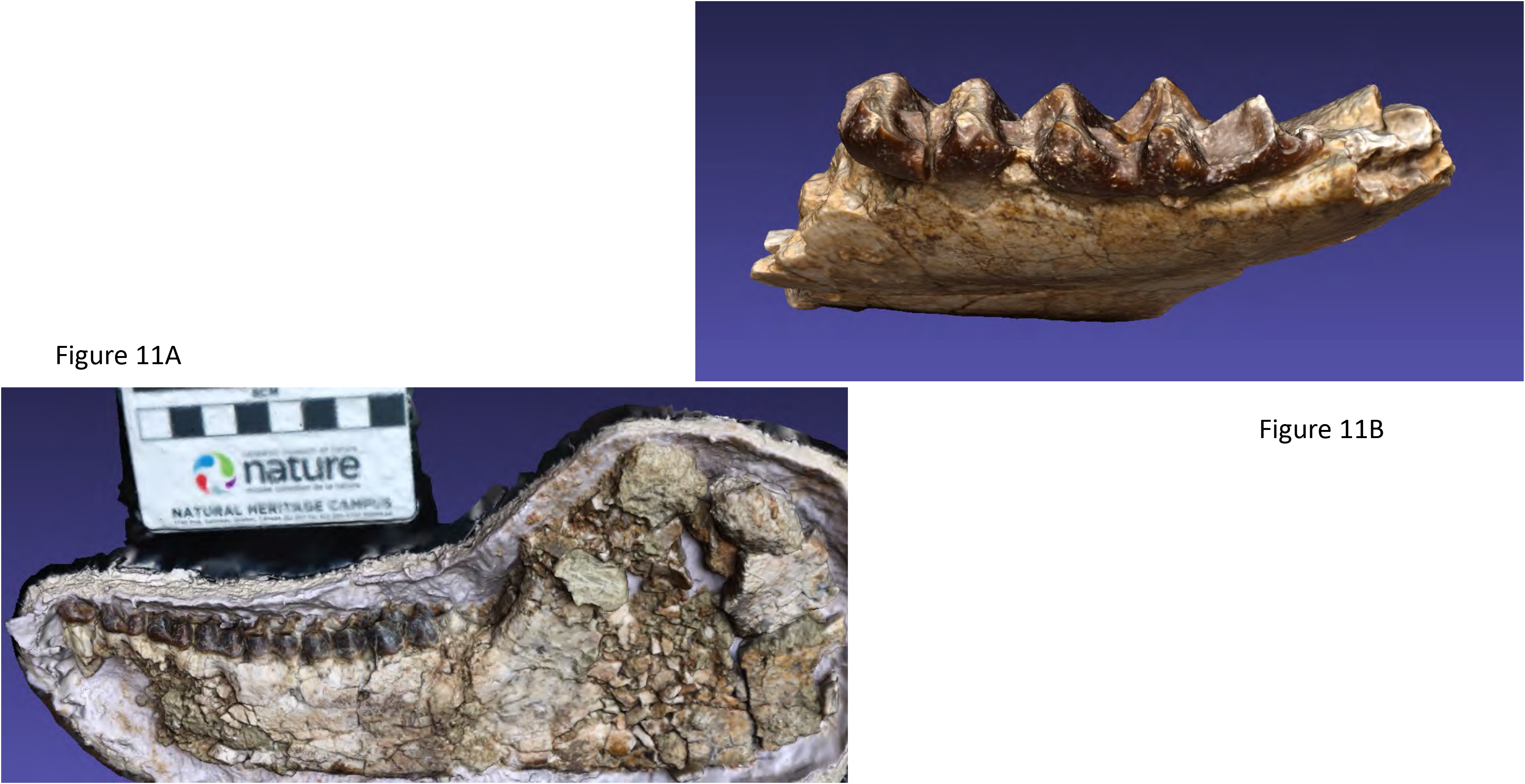
**A)** Hyracoid left mandible GPL2018-P-1. **B)** Hyracoid right mandibular fragment, GPL2018-P-2.

**Table 4.** Measurements of hyracoid teeth in mm.

| **PPG2018-P-1** | Side | mesio-distal | bucco-lingual |
| --- | --- | --- | --- |
| p1 | Lt | 12.81 | n.a. |
| p2 | Lt | 14.64 | n.a. |
| p3 | Lt | 14.96 | n.a. |
| p4 | Lt | 15.72 | n.a. |
| m1 | Lt | 17.59 | n.a. |
| m2 | Lt | 19.61 | n.a. |
| m3 | Lt | 31.01 | n.a. |

| **PPG2018-P-2** |  |  |  |
| --- | --- | --- | --- |
| p3 | Rt | 14.48 | 8.93 |
| p4 | Rt | 16.14 | 10.26 |
| m1 | Rt | 16.63 | 12.60 |
| m2 | Rt | 20.77 | 15.31 |
| m3 | Rt | 32.40 | 14.46 |

To compare the Gorongosa mandibles with those from other sites, we carried out a Principal Component Analysis of dental shape variables. For the left hemimandible (PPG2018-P-1) we used five curves with 15 landmarks each from the buccal side (given that the lingual side is obscured by the plaster jacket) to produce dental row outlines from p3 to m3 (Figure 12A). These landmarks were collected using the software Landmark Editor 3.6 (Wiley et al., 2005). We chose the p3-m3 sequence (excluding p1-p2) to maximize the number of comparative specimens that could be used. We obtained similar outlines from the 3D models of 14 hyracoids. Three of these comparative specimens are housed at the National Museums of Kenya (NMK) and were digitized using photogrammetry following the protocol described by Bucchi and colleagues (Bucchi, Luengo, Fuentes, Arellano-Villalón, & Lorenzo, 2020). Eleven additional comparative specimens were downloaded from Morphosource https://www.morphosource.org/ (Boyer, Gunnell, Kaufman, & McGeary, 2016) (Supplementary Table S7). This comparative sample included the genera *Saghatherium*, *Thyrohyrax*, *Megalohyrax*, *Afrohyrax* and the modern genera *Dendrohyrax* and *Procavia*. The first and last landmarks from each one of the five curves were treated as fixed (i.e., 10 fixed landmarks), whereas all the rest of them (i.e., 65 landmarks) were considered as semi-landmarks. See Supporting Information: Morphometric Analysis: Hyracoidea for further details. This PCA shows that the Gorongosa mandible is closer to specimens of Saghatheriidae (*Saghatherium*, *Thyrohyrax*, and *Megalohyrax*) than to Titanohyracidae (*Afrohyrax*) or modern Procaviidae (*Dendrohyrax* and *Procavia*) (Figure 12B), at least when considering the two first principal components (PCs) that account for ∼70% of the variance of the sample.

**Figure 12.**
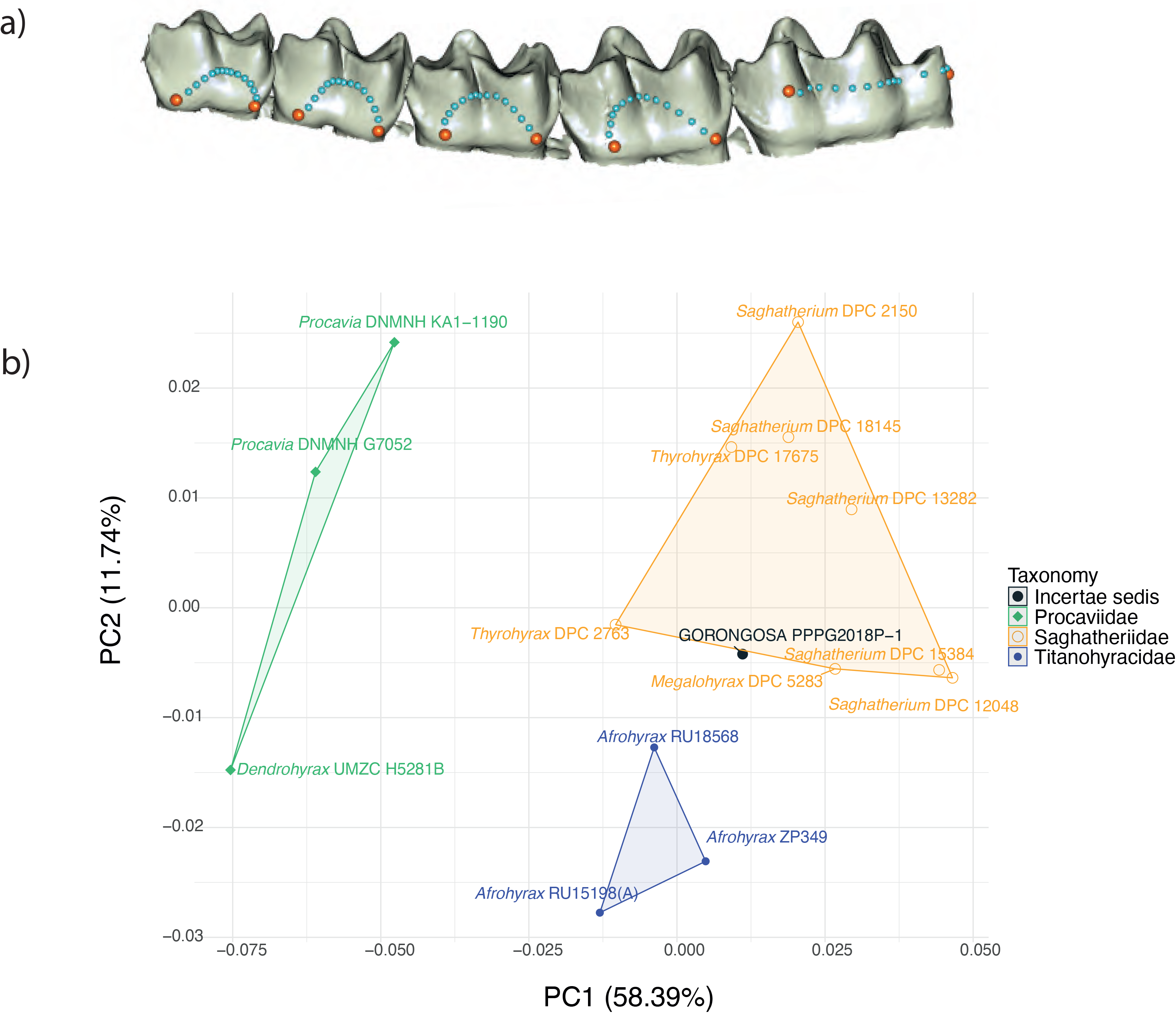
**A)** *Thyrohyrax* specimen (DPC 2763) showing the landmarks (orange spheres) and semi-landmarks (light blue spheres) used in this study. This specimen was selected to display the 3D coordinates as it corresponds to the specimen closest to the multivariate mean in this analysis. **B)** Principal component analysis (PCA) of the dental shape variables (only the two first PCs are shown).

In another analysis using only the m3 from mandible PPG2018-P-2, we used four curves with 10 landmarks each (Figure 13A). This dataset was then compared with the 3D models of 25 hyracoids. Thirteen of these specimens are also housed at the NMK and that were digitized using photogrammetry using the same protocol that was described above, whilst the rest of the sample was obtained from Morphosource https://www.morphosource.org/ (Supplementary Table S8). The comparative sample derives from five families of Hyracoidea: Geniohyidae (*Bunohyrax*), Saghatheriidae (Saghatherium, *Thyrohyrax*, *Megalohyrax*), Titanohyracidae (*Afrohyrax*, *Mereohyrax*), Pliohyracidae (*Parapliohyrax*), and Procaviidae (*Dendrohyrax* and *Procavia*). This dataset was also subjected to a General Procrustes Analysis (GPA) to obtain shape variables. The first and last landmarks from each one of the four curves were treated as fixed (i.e., eight fixed landmarks), whilst the remaining 3D coordinates (i.e., 32 landmarks) were considered as semi-landmark and were slid by using Procrustes distance minimization as criterion. The obtained shape residuals were then used to carry out a PCA. This PCA shows that the Gorongosa m3 is closer to specimens of Saghatheriidae than to those of other families (Figure 13B), at least when considering the first two PCs that account for ∼64% of the variance of the sample. Although most similar to the genus *Megalohyrax*, the Gorongosa specimen does not fully match any of the known taxa within this genus and represents a new species that we are fully describing in a separate manuscript.

Order Primates *inc. sed.* Linnaeus, 1758
Anthropoidea Mivart, 1864
Infraorder Catarrhini Geoffroy, 1812
Referred specimen: PPG2017-P-126

**Figure 13.**
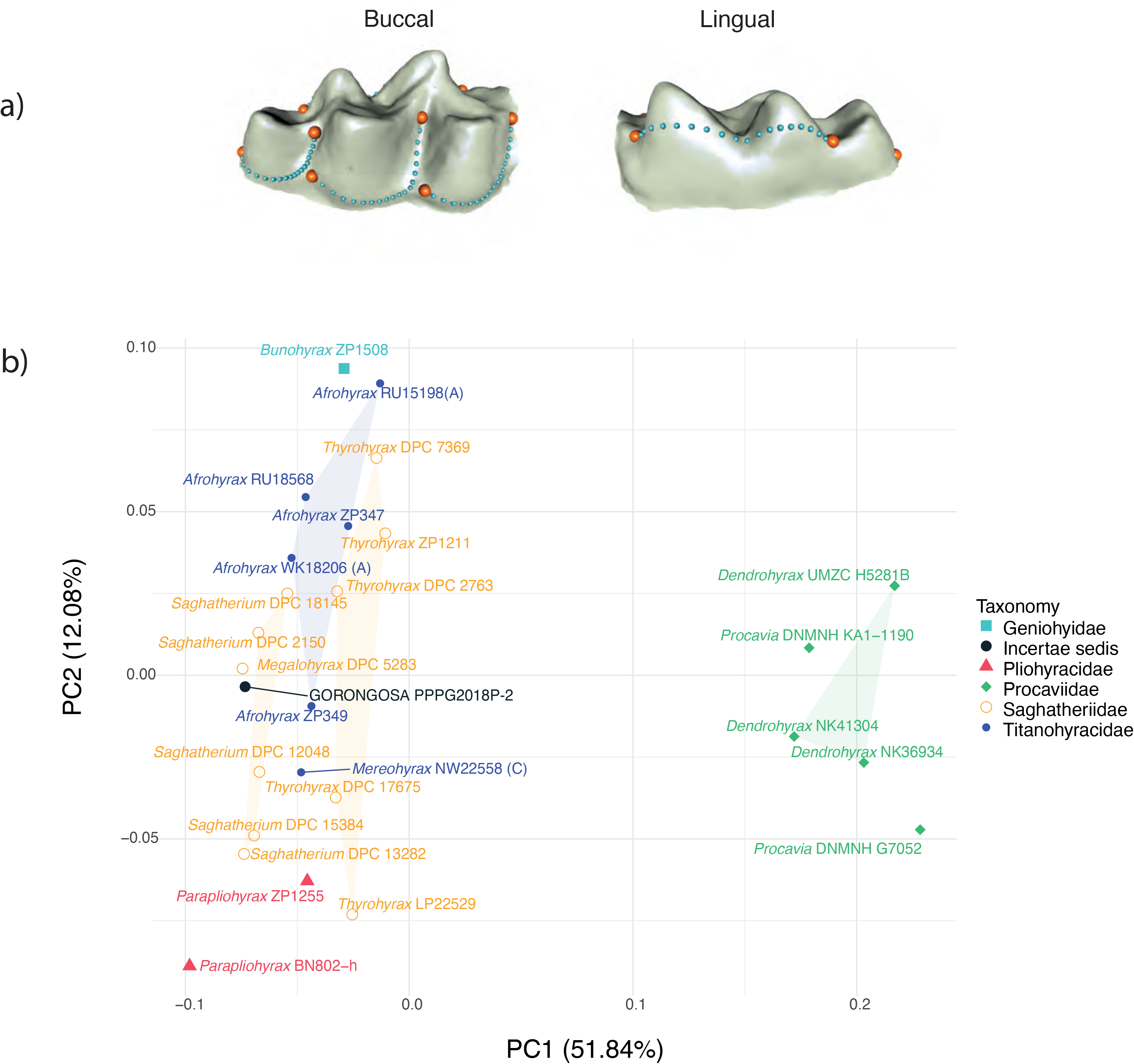
**A)** *Afrohyrax* specimen (ZP349) showing the landmarks (orange spheres) and semi-landmarks (light blue spheres) used in this study. This specimen was selected to display the 3D coordinates as it corresponds to the specimen closest to the multivariate mean in this analysis. **B)** Principal component analysis (PCA) of the m3 shape variables (only the two first PCs are shown).

A single fossil specimen, PPG2017-P-126, was found at GPL-7. This specimen is a left maxillary fragment preserving part of the alveolar process, the malar process, and part of the zygomatic bone (Figure 14). The zygomatic process of the maxilla is broad and horizontal in orientation, unlike the condition usually observed in catarrhine primates. It extends 24 mm from the alveolar process to its lateral end. Laterally the maxilla-zygomatic suture is preserved. The superior aspect of the malar process preserves an area with cortical bone surface that may be the inferior floor of the orbit or an infraorbital canal (Figure 14). In addition, the posterior aspect of this bone is curved, and the cortical bone surface is preserved. Inspection of the microCT scans does not reveal any presence of the floor of a maxillary sinus which, among African catarrhine primates, is present only in hominoids and macaques (Rae & Koppe, 2004). If present, the maxillary sinus would be located above the root apices and could expand into the zygomatic bone. In contrast and worth noting, the cancellous bone of both the alveolar and zygomatic bones is quite dense and compact, while some of the cancellous bone pores around the root apices appear to be filled in with sediment (Figure 15).

**Figure 14.**
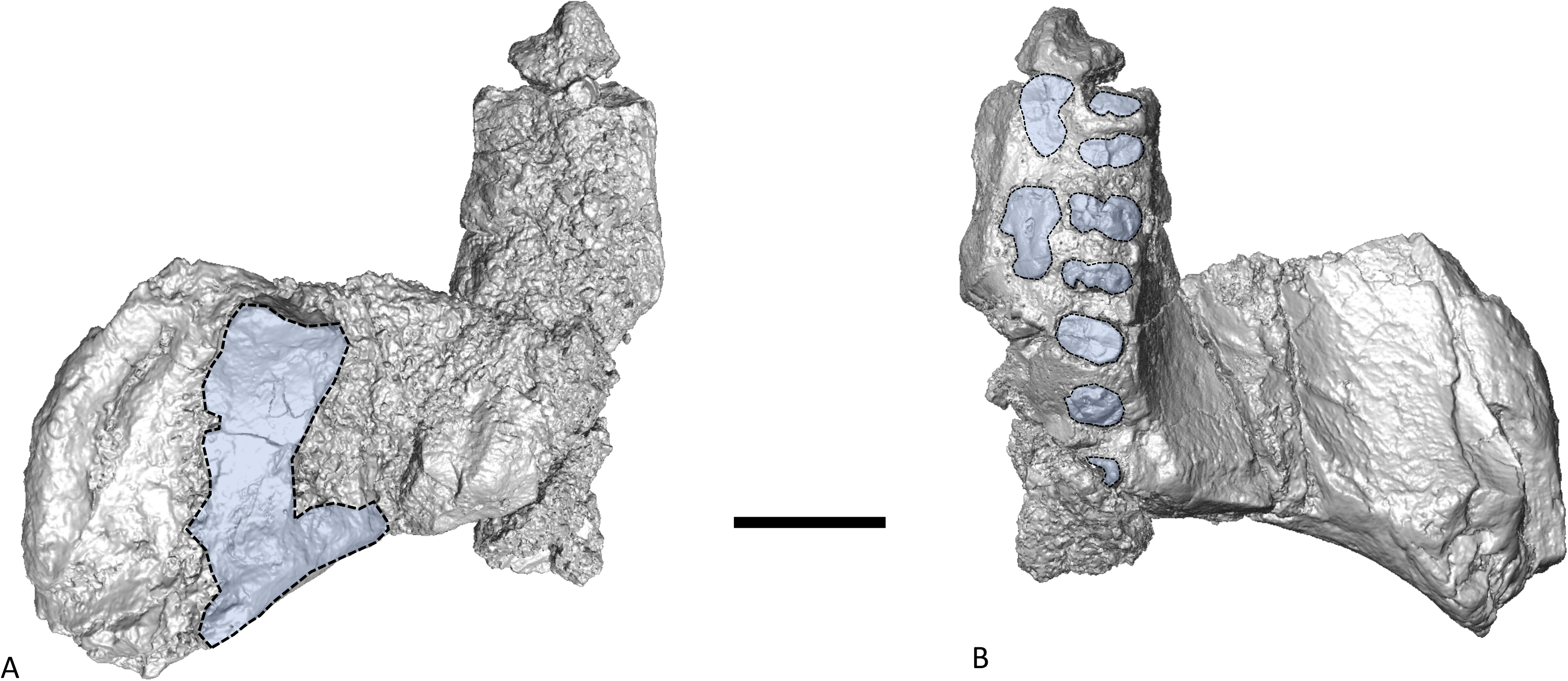
Surface rendering of PPG2017-P-126, left maxillary fragment with alveolar process. **A)** Superior view. In the zygomatic bone some cortical bone surface is still preserved (area highlighted in blue). **B)** inferior view with postcanine tooth roots highlighted in blue. Anterior is to the top. Scale bar = 1 cm.

**Figure 15.**
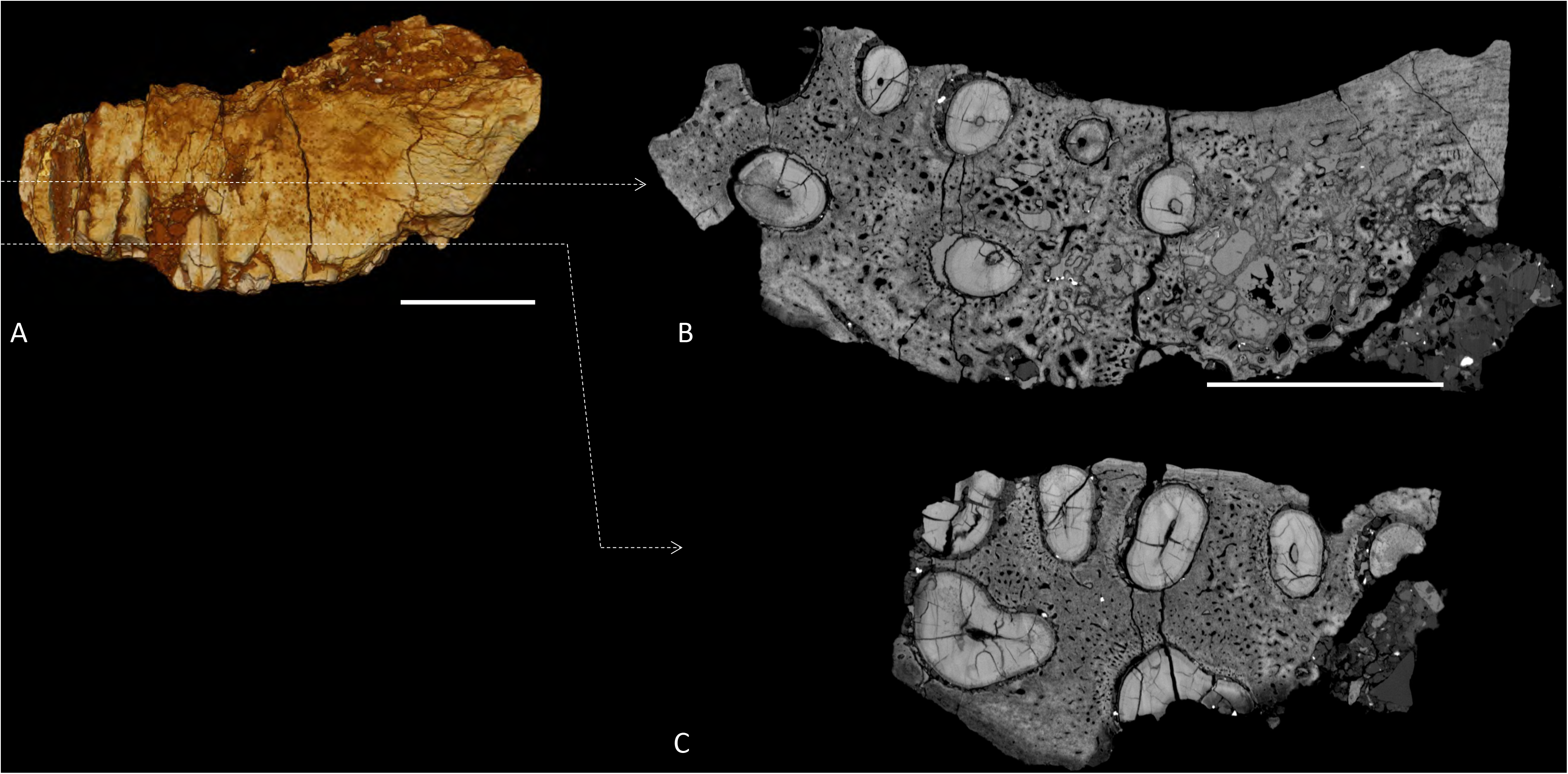
3D rendering and CT sections of PPG2017-P-126, left maxillary fragment with alveolar process. **A)** Buccal view. Anterior to the left. Dashed lines indicate virtual sections at the levels of mid-root and root apex. **B)** Apical view of tooth roots and transverse CT scan showing cancellous bone at the level of root apex. **C)** Apical view of tooth roots and transverse CT scan showing cancellous bone at mid-root. Scale bars = 1 cm.

The tooth crowns are missing, but the alveolar process preserves a series of roots below the furcation, which we interpret as a premolar-molar sequence (Figure 16A and 16B). On the mesial end, a portion of the distal wall of the palatal root of the P3 is still present, while the first set of 3 roots represents a P4, with two buccal roots (the mesio-buccal one is broken) and one palatal root. The second set of 3 roots is M1, again with two buccal roots and one palatal root. The third set of 3 roots is M2, with two buccal roots and a broken palatal root. Distally, there is a small fragment of root encased in sediment that is part of M3. In P4-M2 the lingual root is placed symmetrically between the two buccal roots forming an approximately equilateral triangle. In cross-section it is gutter-shaped along its buccal aspect. The root canals are elliptical in cross-section and relatively small for the dentine present, when compared to a sample of East and South African Miocene and Plio-Pleistocene catarrhine primates. The distal root of M2 has a smaller diameter than the other molar roots. The base of the zygomatic process of the maxilla extends from the P4-M1 junction anteriorly to M3 posteriorly. The mesiodistal length from the mesial root of P4 to the distal root of M2 is 24 mm. The maximum mesiodistal length of M1 at the roots is 8 mm, and the maximum buccolingual width is 9 mm. The overall morphology of the maxillary roots and their cross-sectional shape suggest that the specimen may belong to an anthropoid primate, as noted below.

**Figure 16.**
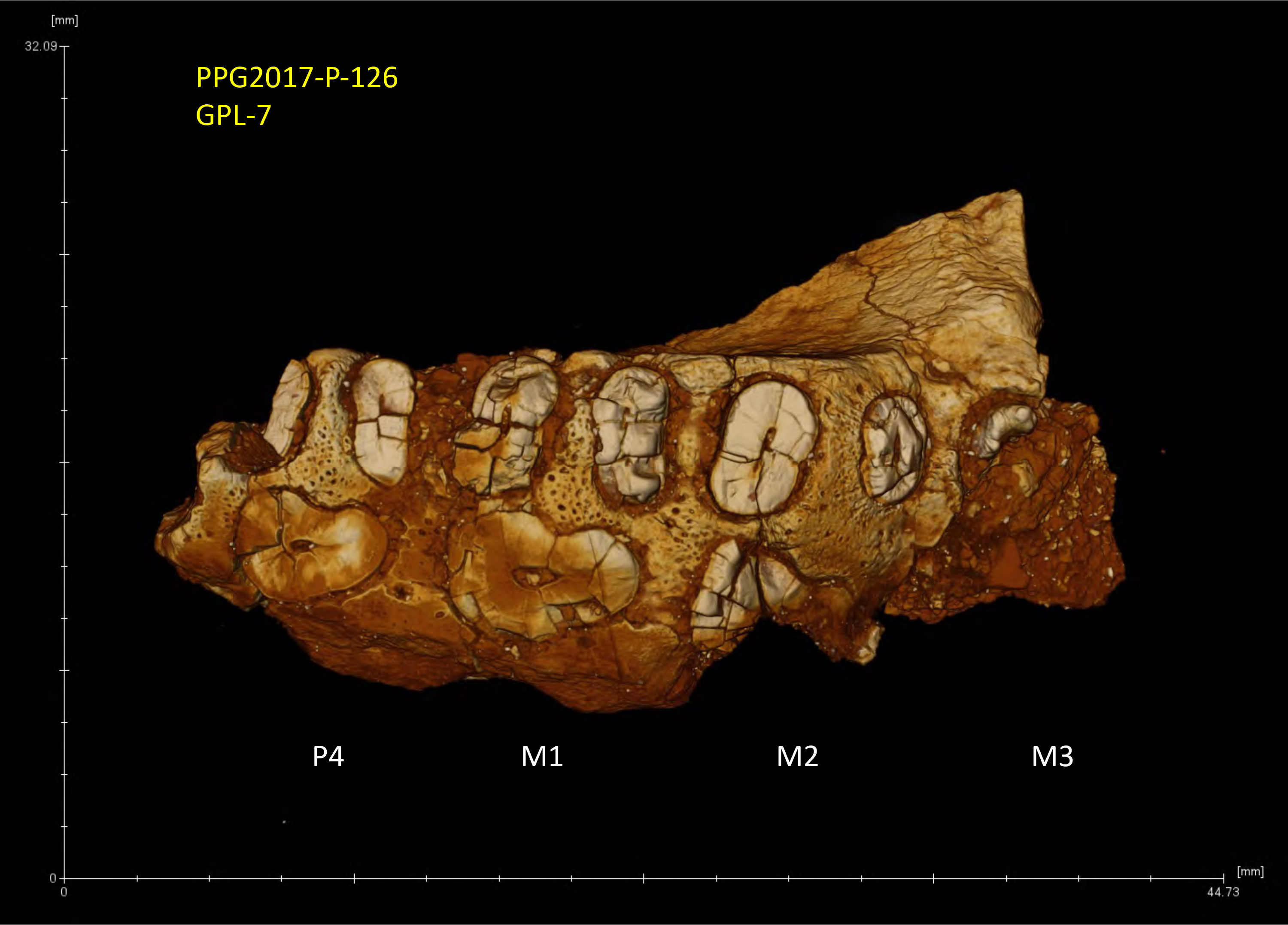
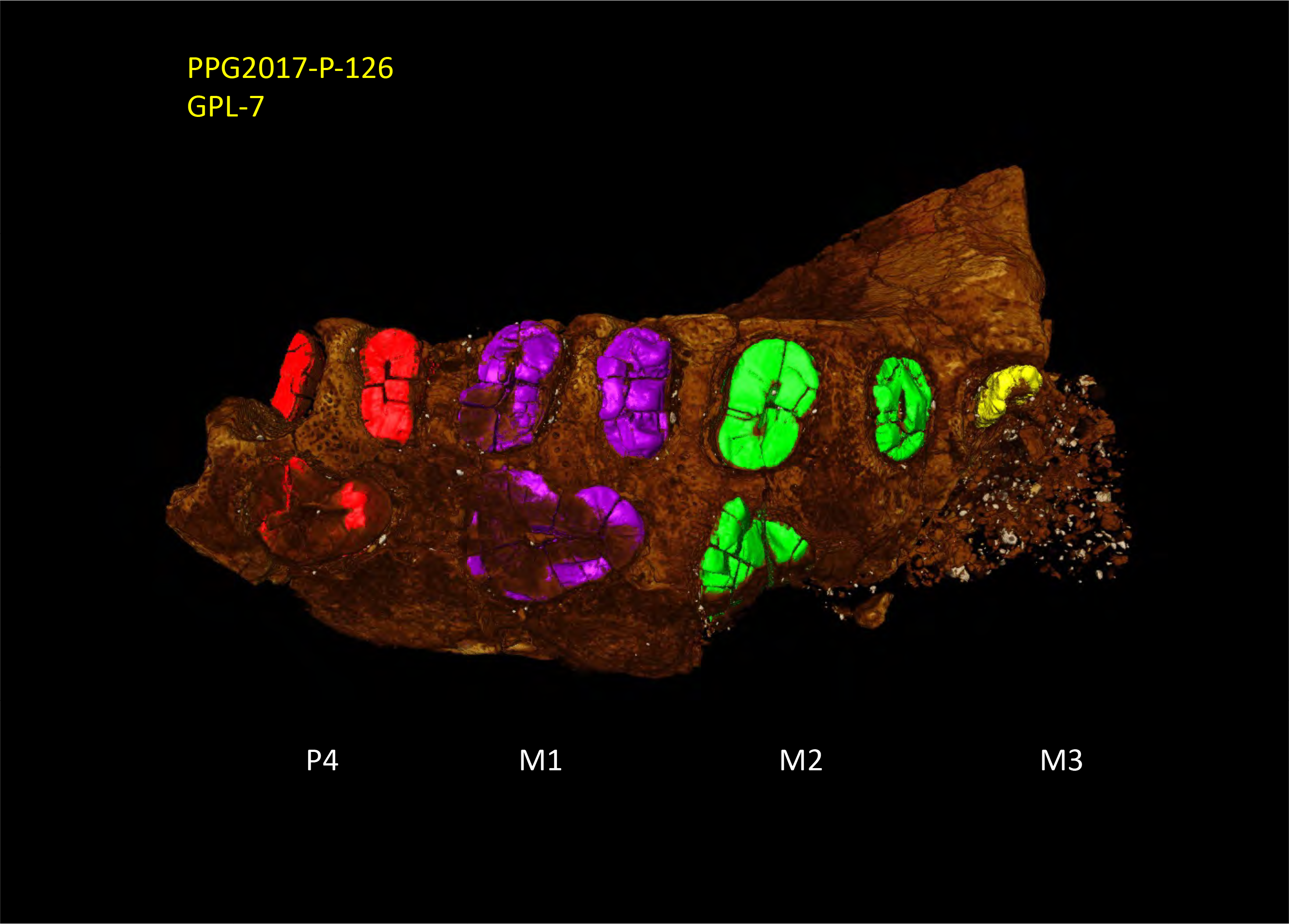
**A)** Inferior view of specimen PPG2017-P-126 showing the broken tooth roots. **B)**Inferior view of specimen PPG2017-P-126 with the tooth roots highlighted in color.

To better assess the morphological affinities of this specimen, we carried out geometric morphometric analyses of the roots. It has been shown that different primate species exhibit distinct tooth root morphologies and that the analysis of tooth root cross-sectional shape is diagnostic both in terms of designating taxonomic status and determining position within the tooth row (Kullmer et al., 2011; Kupczik, 2003; Kupczik & Dean, 2008; Kupczik, Delezene, & Skinner, 2019; Kupczik & Hublin, 2010; Kupczik, Toro-Ibacache, & Macho, 2018). Even though the available morphology is too fragmentary to properly ascribe PPG2017-P-126 to a low taxonomic level, there is enough information to assess its possible primate status, as well as to carry out preliminary phenetic comparisons. Hence, we analyzed the M1 cross-sectional shape, as it corresponds to the most complete set of tooth roots available. We assembled a comparative sample comprising the CT-stacks of several different extinct and extant primates (Supplementary Table S9 and Figure 17). To obtain comparable root outlines, we manually selected the cross-sectional slices that were at the same level as the exposed roots of PPG2017-P-126 for each one the comparative specimens. Then, 30 equidistant 2D semi-landmarks were collected along the outlines of each one of the three roots available for M1, which resulted in a total of 90 2D coordinates (see Supporting Information: Morphometric Analysis: Primates, and Figure 18A). The first landmark of each one of three root outlines (i.e., landmarks 1, 31 and 61) were treated as fixed, and these were defined as the most lingual points for each one of the M1’s root outlines. All the rest of the 2D coordinates (i.e., 87 landmarks) were considered as semi-landmarks. We computed the multivariate mean shape of the sample using the obtained shape variables and the Procrustes distances between this average M1 root shape and each one of the specimens under analysis was calculated (Figure 18B). We plotted all specimens ordered by their Procrustes distance from the mean shape. The median distance (unbroken line) and upper quartile (dashed lines) summarize the distances from the mean shape. The most distinct morphologies are those represented by the papionins present in our sample (*Papio angusticeps* and *Theropithecus oswaldi*). They are the only specimens above the upper quartile in our sample and that could be considered as ‘morphological outliers’ in our analysis. PPG2017-P-126 is close to the median, thus exhibiting an M1 root cross-sectional shape that is not unusual for catarrhine primates. In addition, we used the obtained shape variables to perform a PCA. This PCA shows that the PPG2017-P-126 is closer to *Cercopithecoides williamsi*, *Simiolus enjessi*, and some chimpanzees (Figure 18C), at least when considering the two first principal components (PCs) that account for ∼62% of the variance of the sample. PPG2017-P-126 is located between the Cercopithecoidea and Hominoidea along PC1, whilst along PC2 it shows values like some chimpanzees and some orangutans. Based on these results, we suggest that the root anatomy of PPG2017-P-126 shows primate affinities as observed in species of Catarrhini.

**Figure 17:**
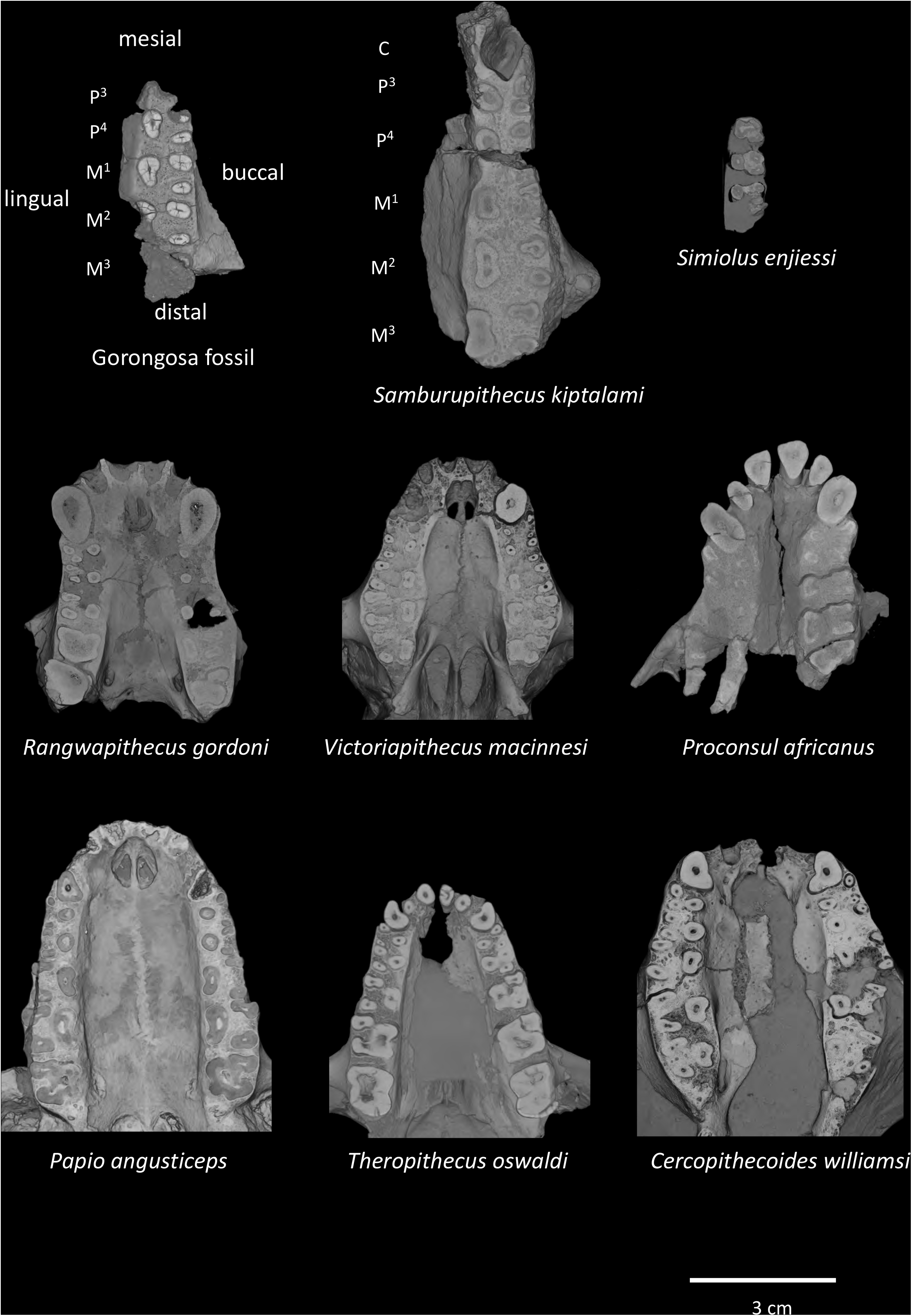
Micro-CT scan of Gorongosa fossil PPG2017-P-126 in inferior view to compare root cross-sections, root canals, and part of the dental arcade with other Miocene and Plio-Pleistocene catarrhine fossil primates from eastern and southern Africa. Anterior is to the top.

**Figure 18.**
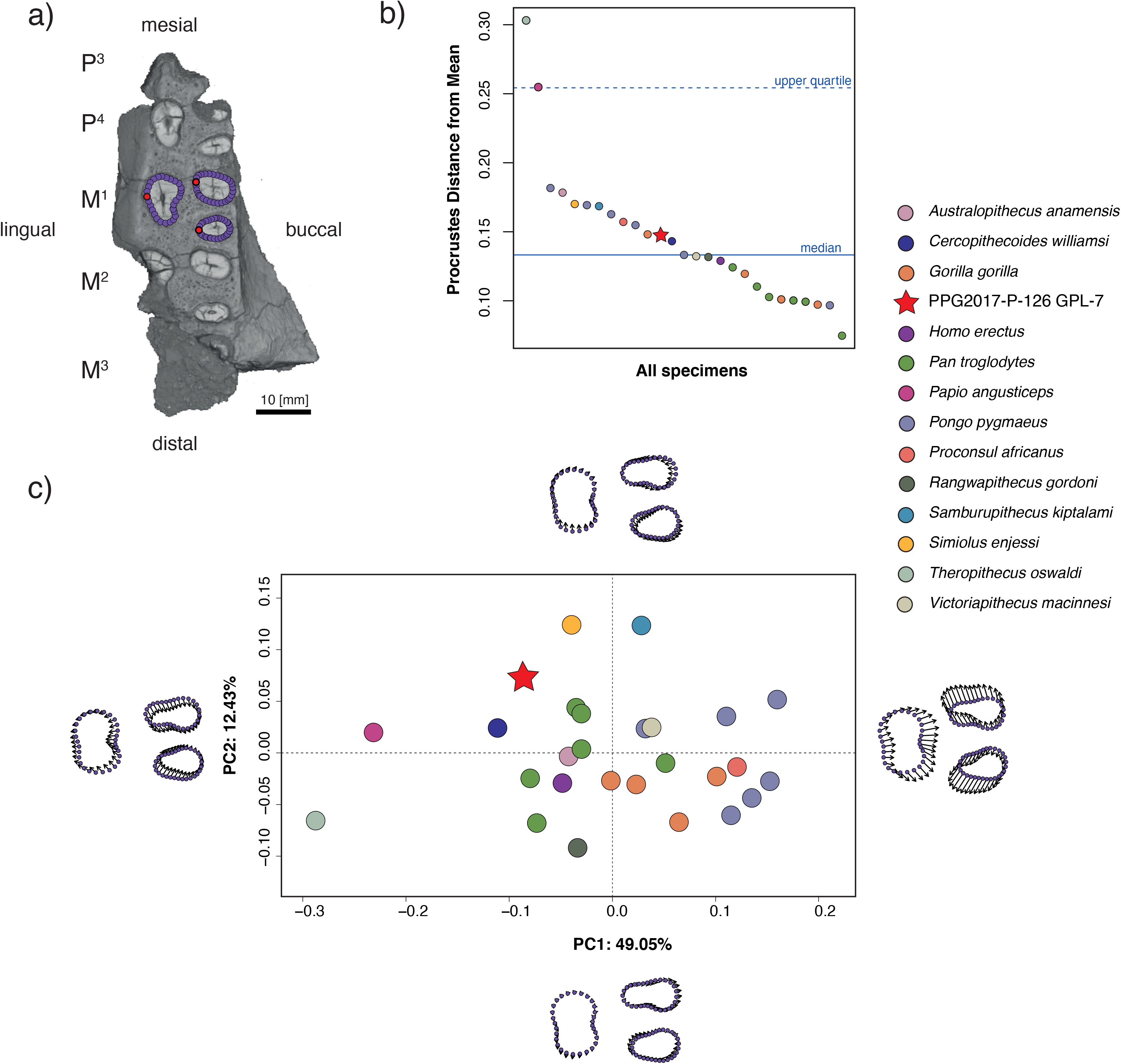
**A)** Gorongosa specimen PPG2017-P-126 showing an outline of the M1 roots cross-section with 30 equidistant landmarks and the most lingual point of each root marked in red. **B)** Multivariate mean shape of the primate specimens using the obtained shape variables and the Procrustes distances between the average M1 root shape and each one of the specimens under analysis. Specimens ordered by their Procrustes distance from the mean shape. The median distance (unbroken line) and upper quartile (dashed line) summarize the distances from the mean shape. **C)** Principal Component Analysis (PCA) of the cross-sectional root shapes showing that the Gorongosa specimen is closest to *Cercopithecoides williamsi*, *Simiolus enjessi*, and some chimpanzees when considering the two first principal components (PCs) that account for ∼62% of the variance of the sample.

## Discussion and conclusions

The new fossil sites from Gorongosa National Park open an entirely new vista on a region of Africa that, until now, had remained paleontologically unknown until now (Figures 1 and 2). No other sites along the East African Rift System yield the combination of fossil woods (e.g., African mahogany), marine invertebrates (crabs, gastropods, bivalves), marine vertebrates (sharks and rays), and terrestrial mammals (e.g., hyracoids). The geological, sedimentological, paleobotanical, geochemical, and paleontological evidence indicates that the Gorongosa fossil sites formed in coastal settings, even though today these sites are ∼95 km from the coast and at ∼100-120 m above sea level (Figures 3 and 4).

The new fossils derive from multiple sedimentary beds across ten paleontological localities in the lower member of the Mazamba Formation. Previous geological work assigned this sedimentary sequence broadly to the Miocene (Real, 1966, Flores, 1973, Tinley, 1976), but no radiometric dates had been obtained prior to our work. Here we have presented the first atmospheric beryllium dates for the Mazamba Formation (Table 4). Atmospheric beryllium samples from the lower member range in age from the early to the late Miocene and confirm the broad placement of this part of the sequence in the Miocene. Two samples from the lowermost sections of GPL-12 provide an early Miocene age for the fluvio-deltaic sediments from which some key fossils derive. Atmospheric beryllium samples from GPL-2, which we expect to be younger based on our tentative correlations (Figures 3 and 4), indicate a late Miocene age for those sediments (Table 4). Thus, the dates obtained so far indicate that the new fossils derive from different intervals of the Miocene, but further applications of different dating techniques are needed to corroborate the chronology suggested by the atmospheric ^10^Be method.

The sedimentological, isotopic, paleobotanical, and paleontological evidence presented here indicates that the fossil sites formed in coastal woodlands/forests or estuarine conditions. At GPL-1, for example, paleosol carbon and oxygen isotopes indicate the prevalence of C_3_ vegetation (trees, shrubs) with some areas of grassland under mesic climate with a high supply of fresh water (Figure 5). This view is supported by the fossil wood (Figure 6), whose most abundant component is *Entandrophragmoxylon* (African mahogany) (Figure 7), a genus that typically grows in areas of high rainfall. There were also palm trees of the genus *Hyphaene*, which are widespread in the humid, hot lowlands with high water tables of tropical Africa today. Other trees in the ancient Gorongosa landscapes include *Terminalioxylon*, which includes some mangrove species, *Zizyphus*, which is common along the edges of watercourses, and *Zanha*, a genus associated with open woodland to dense ravines and riverine forests. Cross sections of the fossil wood vessels indicate the presence of mesophytic trees that cannot tolerate water stress. Thus, these different lines of evidence indicate that terrestrial environments near the coast were consistently warm and wooded, with a prevalence of C_3_ vegetation under mesic conditions.

The rivers descending from the west meandered on a low gradient coastal plain, where they gave rise to estuaries near shallow marine environments (Habermann et al., 2019). Sharks of the genus *Galeocerdo* (Figures 8, 9) were top predators in these estuaries and nearshore environments. Specimens of *Galeocerdo* are known from the Eocene to the present (Türtscher et al., 2021), whilst the species *G. aduncus*, present in the Gorongosa sample, has a temporal range from the Oligocene to the late Miocene (Soto Ovalle, 2016; Türtscher et al., 2021). The genus was widely distributed in the tropical and temperate seas of the Miocene, with specimens found in Madagascar (Andrianavalona et al., 2015), North Africa (Argyriou et al., 2015; Cook, Murray, Simons, Attia, & Chatrath, 2010), Oceania (Fitzgerald, 2004), Eurasia (Marsili, Carnevale, Danese, Bianucci, & Landini, 2007; Villafaña et al., 2020), and the Americas (Carrillo-Briceño, Maxwell, Aguilera, Sánchez, & Sánchez-Villagra, 2015; Landini et al., 2017). Modern *Galeocerdo* ranges from pelagic waters to nearshore environments in tropical and subtropical marine ecosystems, often occurring in river estuaries. Tiger sharks are top predators, with a diet of cephalopods, fish, turtles and other vertebrates (Cortés, 1999). Like the modern tiger sharks, *Galeocerdo* in the past was a highly mobile apex predator that played a major role in structuring coastal ecosystems (Dicken et al., 2017). The presence of these shark fossils in the Miocene of GNP is consistent with our interpretation of estuarine depositional environments (Habermann et al., 2019).

The fossil mammals analyzed here include species of Hyracoidea and Primates. Undoubtedly this is an incomplete list of the taxonomic diversity of Gorongosa fossil mammals, as many of the specimens were only recently excavated and are still under curation and analysis. Hyraxes (order Hyracoidea) belong to the Afrotheria, a clade of mammals with deep evolutionary roots in Africa. There are five species of modern hyraxes, all in the family Procaviidae, but in the past there were at least four additional families: Geniohyidae, Saghatheriidae, Titanohyracidae, and Pliohyracidae. Hyracoids in the Paleogene of Africa were abundant and diverse, both taxonomically and functionally, but declined in overall diversity during the late Miocene (Rasmussen & Gutiérrez, 2010). The chewing teeth of the Gorongosa hyracoid are brachydont and bilophodont, very likely for a diet of relatively soft leaves. The Gorongosa hyracoids represent a very large species with affinities to taxa in the family Saghatheriidae, but different from currently known species (Figures 11, 12, 13). The family Saghatheriidae includes the genera *Microhyrax*, *Saghatherium*, *Selenohyrax*, *Thyrohyrax*, *Megalohyrax*, and *Regubahyrax* spanning from the Eocene to the early Miocene. Specimens of *Regubahyrax* from the early Miocene of Libya document the latest known occurrence of saghatheriids (Pickford, 2009). The lower molars of *Regubahyrax* have well-developed cristids and spurs (Pickford, 2009), but the spurs are not developed in the Gorongosa hyracoid. Although most similar to the genus *Megalohyrax*, the Gorongosa specimen does not fully match any of the known taxa within this genus and represents a new species that we are fully describing in a separate manuscript (Bobe et al. in preparation).

Catarrhine primates also occurred in the ancient Gorongosa landscapes (Figures 14-18). The available morphology of the Gorongosa specimen (PPG2017-P-126) is too fragmentary to ascribe it to a low taxonomic level, but our comparisons of the molar root cross-sectional shape show that PPG2017-P-126 is not unusual for catarrhine primates. Our analyses of the Gorongosa specimen place it close to the primates *Cercopithecoides williamsi*, *Simiolus enjessi*, and some chimpanzees. Thus, conservatively, we assign this specimen to an indeterminate species of catarrhine. Paleogene and early Neogene African primates are associated with wooded or forested environments.

The fossils documented here represent the first descriptions of a substantial fossil record that is just emerging. The Gorongosa paleontological record opens up the possibility of testing important hypotheses about the role of the eastern coastal forests in shaping the evolution of African mammals (Joordens et al., 2019; Kingdon, 2003). As the fossil record from Gorongosa is further described and analyzed, it will yield a potent database spanning different intervals of the Miocene, which will then be compared to other sites on the continent. Thus we will be able to assess the effects of the northeast-southwest arid corridor in promoting the geographic isolation and evolutionary trajectories of coastal forest plant and animal communities in the past (Morley & Kingdon, 2013). The Gorongosa fossil record points to the persistence of woodlands and wooded grasslands along the southeastern coast of Africa during the Miocene, but further work is needed to assess the taxonomic affinities of the Gorongosa mammals with contemporaneous faunas elsewhere in Africa.

After four field seasons (2016-2019), extensive surveys, and new approaches in the search of paleontological sites (d’Oliveira Coelho et al., 2021), the Paleo-Primate Project Gorongosa has 1) documented ten new paleontological localities, 2) established a preliminary stratigraphic and sedimentological framework for the fossil sites (Habermann et al., 2019), 3) provided the first radiometric age determinations for the Mazamba Formation, 4) provided the first reconstructions of past vegetation in the region based on pedogenic carbonates and fossil wood, and 5) described the first fossil teeth from the southern East African Rift System. The Gorongosa sample includes new species of fossil mammals, and a unique combination of fossil specimens straddling the terrestrial/marine biomes. The Gorongosa fossil sites provide the first evidence of persistent woodlands and forests on the coastal margins of southeastern Africa during the Miocene.

## Acknowledgements

We are deeply thankful to Greg Carr for his visionary approach to preserving the Gorongosa ecosystem, and for vital support to the Paleo-Primate Project Gorongosa. Research permits were granted by the Direcção Nacional do Património Cultural, Mozambique, with the support of professors Hilário Madiquida and Solange Macamo of Eduardo Mondlane University. We are very grateful to Gorongosa National Park, the Park Warden Pedro Muagura, the *fiscais* who provide support, and the Park staff, including Vasco Galante and Patricia Álvares da Guerra. We appreciate the help with logistics provided by Jason Denlinger and Tongai Castigo. We thank Katarina Almeida-Warren, Robert Anemone, Ana Gledis da Conceição, Gabriela Curtiz, Celina Dias, Katherine Elmes, Roberto Mussibora, Inês Sevene, and students from the Oxford-Gorongosa Paleo-Primate Field School who took part in the excavations and other aspects of field research. Pepson Makanela prepared and curated specimens while providing training to Mozambican students. We thank S. Adnet, B. Beatty, R. Bernor, C. Brochu, D. Domning, M. Fortelius, D. Geraads, F. Guillocheau, T. Harrison, J. Hutchinson, C. Peters, M. Pickford, C. Robin, C. Robinson, K. Stewart, and W. Sanders for insightful discussions about PPPG research. For access to CT scans of and scanning assistance with the comparative primate fossil specimens we thank the Department of Earth Science, National Museums of Kenya, J.-J. Hublin (Max Planck Institute for Evolutionary Anthropology, Leipzig) E. Mbua, F. Spoor, M. Skinner and H. Temming. For access to the tomographic scan data of South African primate fossil specimens via MorphoSource, we thank Ditsong National Museum of Natural History and J. Adams of the Department of Anatomy and Developmental Biology, Monash University. L. Léanni assisted with chemical treatments and ICP-OES measurements in the LN2C laboratories (Cosmogenic Nuclides French National Laboratory, CEREGE, Aix-en-Provence). The ^10^Be and ^26^Al measurements were performed at the ASTER AMS national facility (CEREGE) which is supported by the INSU/CNRS, the ANR through the “Projets thématiques d’excellence” program for the “Equipements d’excellence” ASTER-CEREGE action, IRD. We thank the National Geographic Society for grants to S. Carvalho (NGS-57285R), R. Bobe (NGS-51140R-18), T. Lüdecke and J. Habermann (NGS-51478R). M. Bamford acknowledges funding from PAST (Palaeontological Scientific Trust, South Africa). T. Püschel was funded by the Leverhulme Trust Early Career Fellowship, ECF-2018-264. J. d’Oliveira Coelho was funded by the Portuguese Foundation for Science and Technology (FCT) Grant SFRH/BD/122306/2016.

## SUPPLEMENTARY FIGURE

**Supplementary Figure S1:**
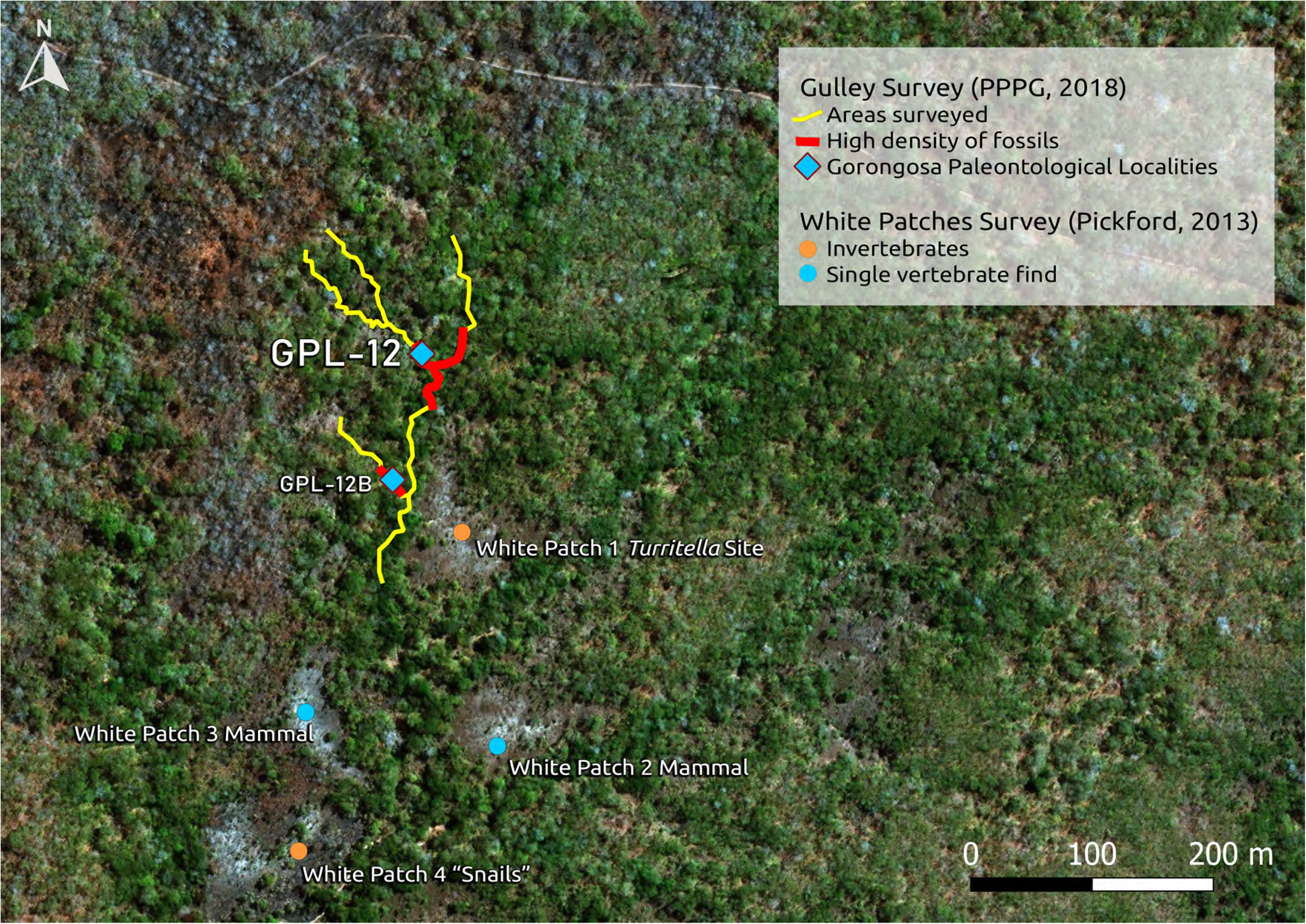
Detail of the location of Gorongosa Paleontological Locality 12 (GPL-12) with a high density of fossils. See also Figure 3C.

## SUPPLEMENTARY TABLES

**Supplementary Table S1:**
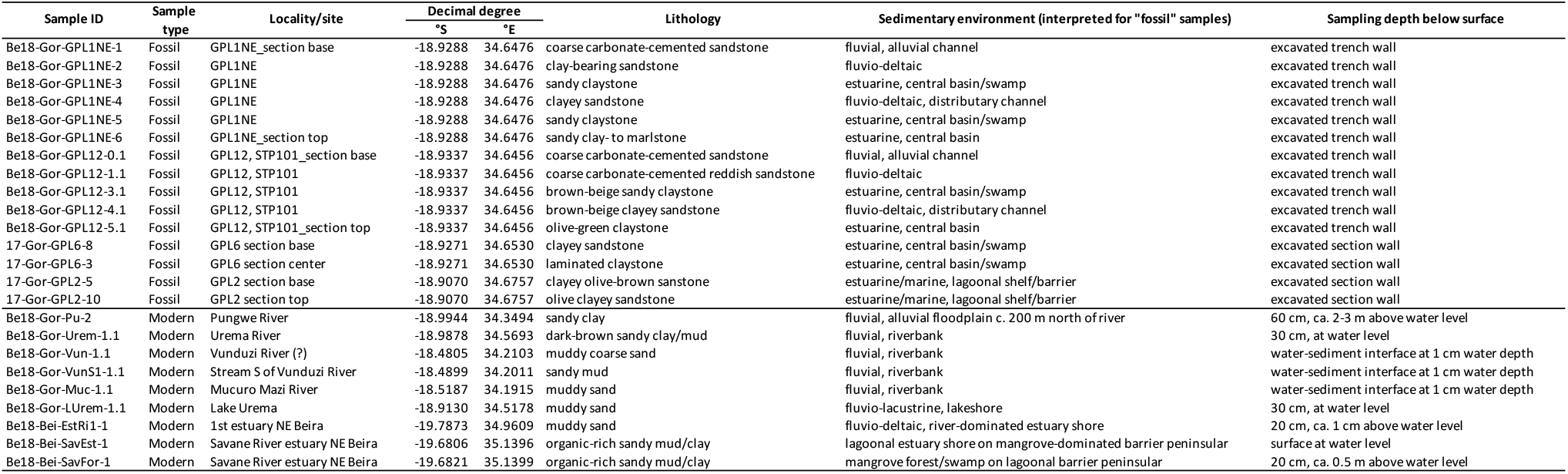
Sampling details for rock samples from the lower member of the Mazamba Formation and modern sediment samples taken for atmospheric ^10^Be dating.

**Supplementary Table S2:** ^10^Be and ^9^Be concentrations and ^10^Be/^9^Be ratios for modern reference samples and fossil samples from the lower member of the Mazamba Formation selected for atmospheric ^10^Be dating. The modern samples and corresponding chemical blank (measured in February 2019) were spiked using 300 µl of a 10^-3^ g.g^-1 9^Be solution (Sharlau 1000 mg/L Be standard), while the lower Mazamba Formation samples and corresponding chemical blank (measured in May 2019) were spiked with ∼150 µl of the LN2C in-house phenakite 3.10^-3^ g.g^-1 9^Be carrier solution. The ^10^Be/^9^Be ratios measured at ASTER were corrected from chemical blank ratios of 1.51 x 10^-14^ for the modern samples and 1.45 x 10^-15^ for the lower Mazamba Formation samples.

| Samples | | Depth<br>[m] | Sample weight<br>[g] | Measured ($^{10}\text{Be}/^9\text{Be}$)<br>* $10^{-13}$ | Authigenic $^9\text{Be}$<br>* $10^{16}$ [at.g$^{-1}$] | Authigenic $^{10}\text{Be}$<br>* $10^7$ [at.g$^{-1}$] | Authigenic $^{10}\text{Be}/^9\text{Be}$<br>* $10^{-8}$ |
| --- | --- | --- | --- | --- | --- | --- | --- |
| Modern | Be18-Gor-Pu-2 | | 0,9593 | 106,6344 ± 2,1964 | 5,1161 ± 0,1496 | 22,6219 ± 0,4653 | 4,4217 ± 0,3162 |
| | Be18-Gor-Urem-1.1 | | 0,9584 | 89,1654 ± 1,9579 | 7,9130 ± 0,2405 | 18,9476 ± 0,4153 | 2,3945 ± 0,1795 |
| | Be18-Gor-Vun-1.1 | | 0,9596 | 15,8407 ± 0,4884 | 1,6814 ± 0,0095 | 3,3464 ± 0,1021 | 1,9902 ± 0,1235 |
| | Be18-Gor-VunS1-1.1 | | 0,9574 | 86,4330 ± 1,8054 | 7,5941 ± 0,0724 | 18,3644 ± 0,3829 | 2,4182 ± 0,1109 |
| | Be18-Gor-Muc-1.1 | | 0,9585 | 84,7570 ± 2,1050 | 2,6621 ± 0,1283 | 17,9818 ± 0,4458 | 6,7548 ± 0,7323 |
| | Be18-Gor-Lurem-1.1 | | 0,9624 | 70,8797 ± 1,7002 | 3,3840 ± 0,1200 | 14,9327 ± 0,3574 | 4,4127 ± 0,3776 |
| | Be18-Bei-EstRi1-1 | | 0,9585 | 280,9860 ± 5,0466 | 4,2827 ± 0,0239 | 59,3875 ± 1,0660 | 13,8668 ± 0,5213 |
| | Be18-Bei-SavEst-1 | | 0,9583 | 77,6644 ± 1,8204 | 2,3125 ± 0,0795 | 16,4663 ± 0,3852 | 7,1206 ± 0,5920 |
| | Be18-Bei-SavFor-1 | | 0,9583 | 75,6599 ± 1,9046 | 2,6573 ± 0,0655 | 16,0133 ± 0,4023 | 6,0262 ± 0,4240 |
| Lower Mazamba Formation | Be18-Gor-GPL1NE-1 (2) | | 0,9162 | 1,7757 ± 0,0005 | 0,0137 ± 0,0569 | 0,5941 ± 0,0186 | 0,1894 ± 0,0137 |
| | Be18-Gor-GPL1NE-2 | | 0,9174 | 0,1918 ± 0,0008 | 0,0011 ± 0,1369 | 0,0610 ± 0,0032 | 0,0096 ± 0,0011 |
| | Be18-Gor-GPL1NE-3 | | 0,9137 | 1,2693 ± 0,0005 | 0,0054 ± 0,1006 | 0,4189 ± 0,0137 | 0,0728 ± 0,0054 |
| | Be18-Gor-GPL1NE-4 (2) | | 0,9158 | 0,0383 ± 0,0018 | 0,0011 ± 0,0457 | 0,0125 ± 0,0015 | 0,0045 ± 0,0011 |
| | Be18-Gor-GPL1NE-5 | | 0,9172 | 0,0082 ± 0,0024 | 0,0003 ± 0,0490 | 0,0027 ± 0,0005 | 0,0009 ± 0,0003 |
| | Be18-Gor-GPL1NE-6 (2) | | 0,9151 | 2,6210 ± 0,0004 | 0,0196 ± 0,1535 | 0,8776 ± 0,0272 | 0,2065 ± 0,0196 |
| | 17-Gor-GPL2-5 | | 0,9167 | 1,5327 ± 0,0005 | 0,0048 ± 0,0700 | 0,5178 ± 0,0162 | 0,0734 ± 0,0048 |
| | 17-Gor-GPL2-10 (2) | | 0,9174 | 1,9923 ± 0,0006 | 5,0197 ± 0,1192 | 0,6586 ± 0,0203 | 0,1312 ± 0,0102 |
| | 17-Gor-GPL6-3 | | 0,9173 | 0,4146 ± 0,0006 | 0,0023 ± 0,0873 | 0,1396 ± 0,0056 | 0,0269 ± 0,0023 |
| | 17-Gor-GPL6-8 | | 0,9192 | 0,1950 ± 0,0009 | 0,0040 ± 0,0568 | 0,0653 ± 0,0041 | 0,0296 ± 0,0040 |
| | Be18-Gor-GPL12-0.1 (2) | | 0,9174 | 0,0464 ± 0,0015 | 0,0006 ± 0,1541 | 0,0156 ± 0,0016 | 0,0027 ± 0,0006 |
| | Be18-Gor-GPL12-1.1 | | 0,9174 | 0,0103 ± 0,0030 | 0,0003 ± 0,0754 | 0,0035 ± 0,0007 | 0,0008 ± 0,0003 |
| | Be18-Gor-GPL12-3.1 (2) | | 0,9175 | 0,4907 ± 0,0006 | 0,0025 ± 0,0883 | 0,1649 ± 0,0071 | 0,0277 ± 0,0025 |
| | Be18-Gor-GPL12-4.1 | | 0,9191 | 0,0423 ± 0,0019 | 0,0008 ± 0,1172 | 0,0141 ± 0,0019 | 0,0030 ± 0,0008 |
| | Be18-Gor-GPL12-5.1 (2) | | 0,9168 | 0,1879 ± 0,0009 | 0,0011 ± 0,0901 | 0,0630 ± 0,0037 | 0,0089 ± 0,0011 |

**Supplementary Table S3.** (in-situ): Results of ^26^Al/^10^Be analyses. Uncertainties (±1σ) include only analytical uncertainties. To each sample, ∼150 µl of the LN2C in-house phenakite 3.10^-3^ g.g^-1 9^Be carrier solution was added. ^27^Al natural concentrations were measured by ICP-OES, and these concentrations were sufficient to perform measurements without addition of an aluminum carrier. The concentration measurements were corrected for the chemical blank ratios of 1.91 ± 0.53 x 10^-15^ and 1.32 ± 0.66 x 10^-15^ for ^10^Be/^9^Be and ^26^Al/^27^Al ratios, respectively.

| Sample | Depth (cm) | Depth<br>(g.cm<sup>-2</sup>) | Dissolved<br>quartz (g) | $^9\text{Be}$ carrier (10<sup>19</sup><br>at.) | $^{10}\text{Be}$ (10<sup>5</sup> at.g<sup>-1</sup>) | $^{26}\text{Al}$ (10<sup>5</sup> at.g<sup>-1</sup>) | $^{26}\text{Al}/^{10}\text{Be}$ |
| --- | --- | --- | --- | --- | --- | --- | --- |
| 16-Gor-Muss-7 | 1500 | 3750 | 20,1345 | 3,0244 | 9909,71 ± 1328,88 | 37881,44 ± 14645,17 | 3,8227 ± 1,5642 |
| 16-Gor-Muss-8 | 1050 | 2625 | 20,1432 | 3,0511 | 11973,11 ± 1589,53 | 57683,13 ± 13054,15 | 4,8177 ± 1,2640 |

**Supplementary Table S4.** (in-situ): Model outputs of burial durations and denudation rates. The data were obtained using a Monte Carlo Regression Model (Braucher, Merchel, Borgomano, & Bourlès, 2011; Pappu et al., 2011). Burial age (expressed here in ka = 1000 a) and denudation rate (expressed in m.Ma^-1^= meter per million years) uncertainties (reported as 1σ) are a propagation of the half-life uncertainties. Parameters used for the calculation: latitude: 19,09°; altitude: 166 m; pressure: 993.5 mbar; mean density: 2.5 g.cm^-3^; Stone Scaling: 0.75; τ^10^Be: 1.387 ± 0.0120 Ma (Chmeleff, von Blanckenburg, Kossert, & Jakob, 2010; Korschinek et al., 2010); τ^26^Al: 0.705 ± 0.024 Ma (Nishiizumi, 2004; Norris, Gancarz, Rokop, & Thomas, 1983); P10 SLHL: 4.03 ± 0.18 at.g^-1^.a^-1^ (Borchers et al., 2016; Molliex et al., 2013); ^10^Be sea level slow muon-induced production: 0.013 ± 0.012 at.g^-1^.a^-1^; ^10^Be sea level fast muon induced production: 0.040 ± 0.004 at.g^-1^.a^-1^; ^26^Al sea level slow muon-induced production: 0.84 ± 0.017 at.g^-1^.a^-1^; ^26^Al sea level fast muon-induced production: 0.081 ± 0.051 at.g^-1^.a^-1^; ^26^Al/^10^Be spallogenic production ratio: 6.61 ± 0.52; Attenuation Length neutrons: 160 g.cm^-2^; Attenuation Length slow muons: 1500 g.cm^-2^; Attenuation Length fast muons: 4320 g.cm^-2^ (Braucher et al., 2011). The studied site’s scaled neutronic production is 3.04 at.g^-1^.a^-1^ for ^10^Be and 20.11 at.g^-1^.a^-1^ for ^26^Al, slow muons production is 0.01 at.g^-1^.a^-1^ for ^10^Be and 0.91 at.g^-1^.a^-1^ for ^26^Al, and fast muons production is 0.04 at.g^-1^.a^-1^ for ^10^Be and 0.08 at.g^-1^.a^-1^ for ^26^Al (Stone, 2000; Braucher et al., 2011). B. = burial; Denud. = denudation; Min = minimum; Max = maximum.

| Sample | Model Without Post-B production |  | Model With Post-B. production |  |  |  |  |
| --- | --- | --- | --- | --- | --- | --- | --- |
|  | Denud. before burial (m.Ma<sup>-1</sup>) | Min Burial duration (ka) | Denud. before B. (m.Ma<sup>-1</sup>) | Max Burial duration (ka) | Denud. after B. (m.Ma<sup>-1</sup>) | % [<sup>10</sup>Be] Post-B. | % [<sup>26</sup>Al] Post-B. |
| 16-Gor-Muss-7 | 140,04 | 1 316,25 ± 539,66 | 1 054,85 | 971,99 ± 398,52 | 20,93 | 84 | 81 |
| 16-Gor-Muss-8 | 147,30 | 838,16 ± 220,96 | 1 746,90 | 971,99 ± 256,24 | 20,93 | 92 | 92 |

**Supplementary Table S5.**
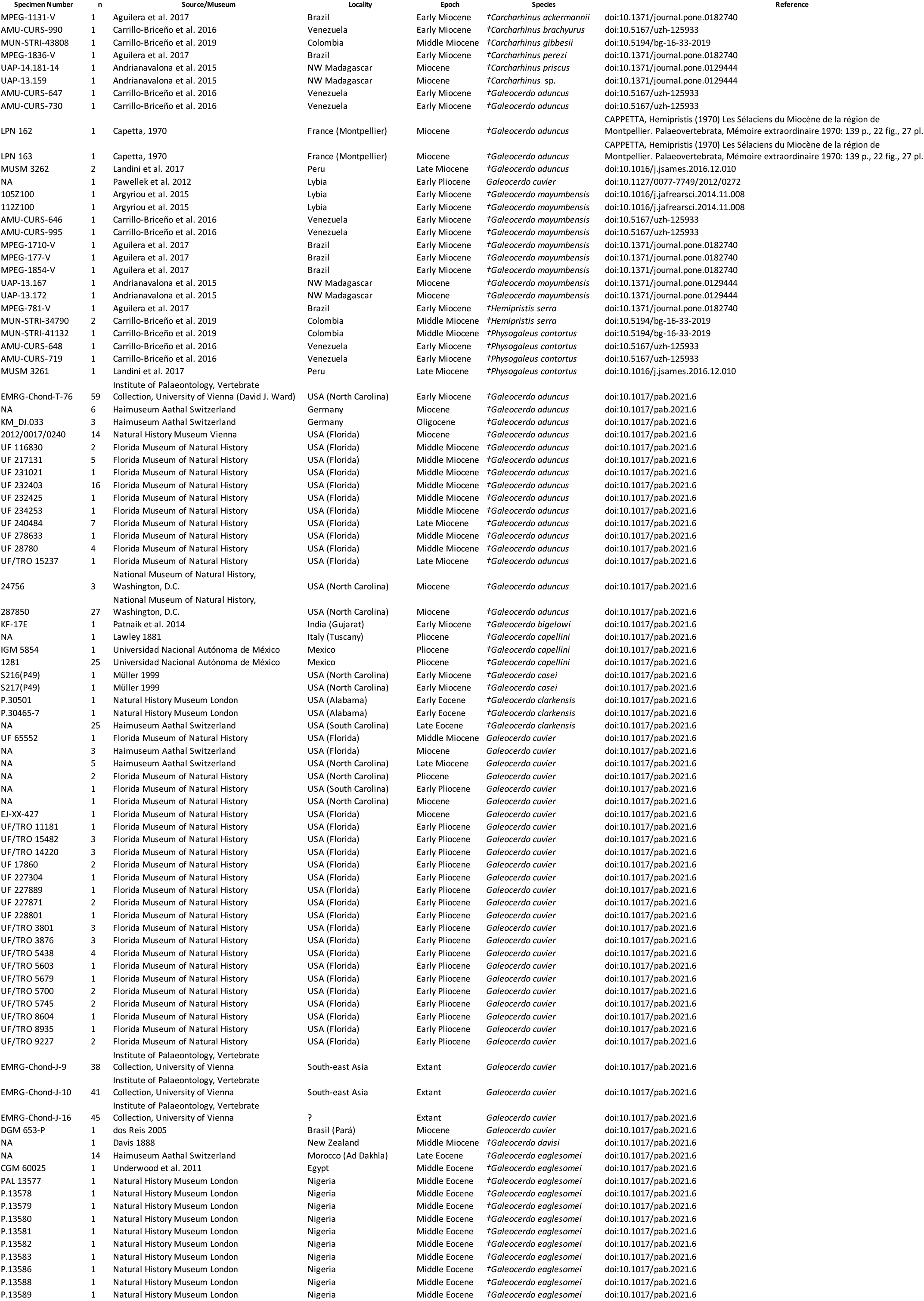
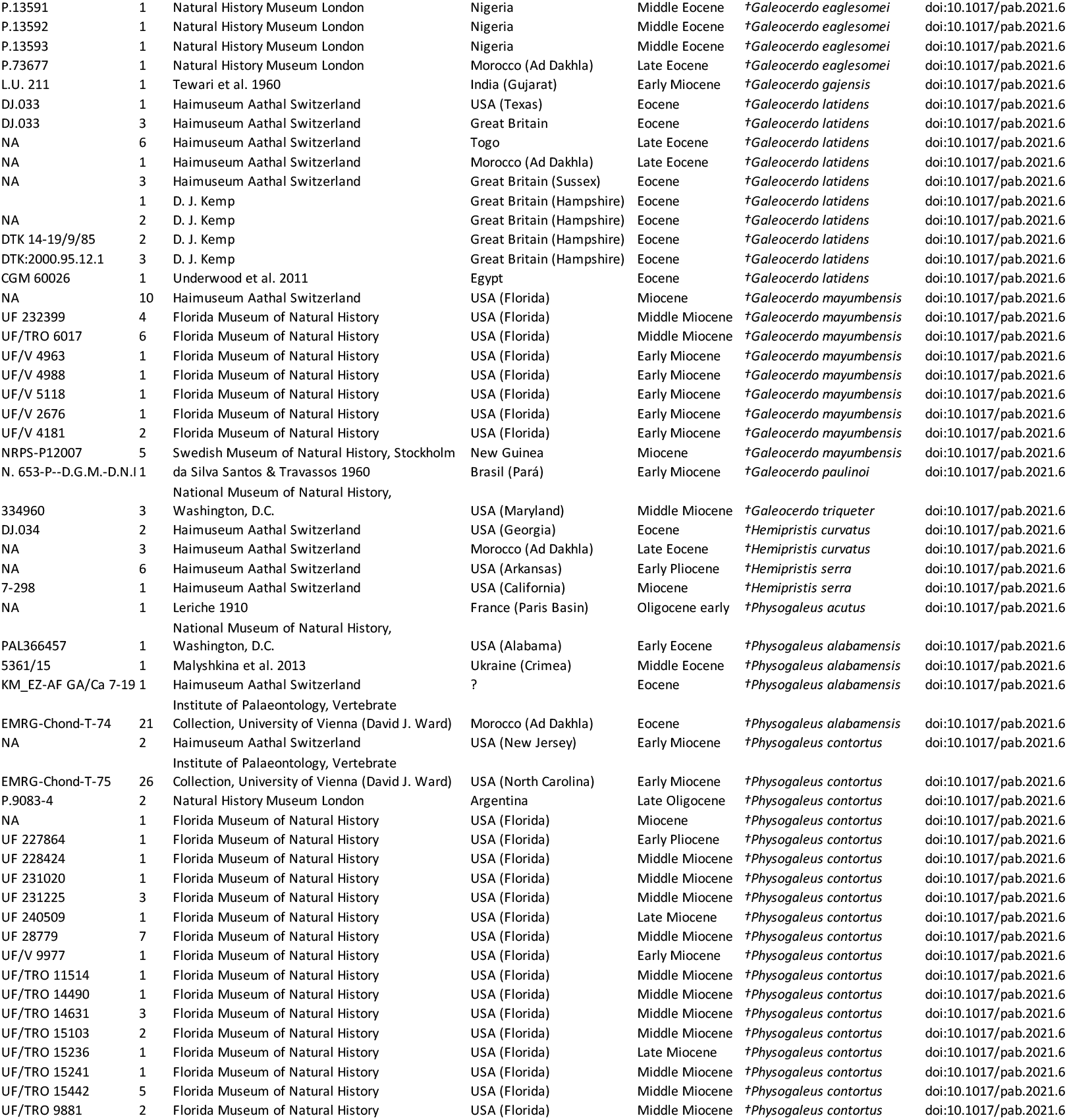
Comparative sample of fossil shark teeth from the published record.

**Supplementary Table S6.**
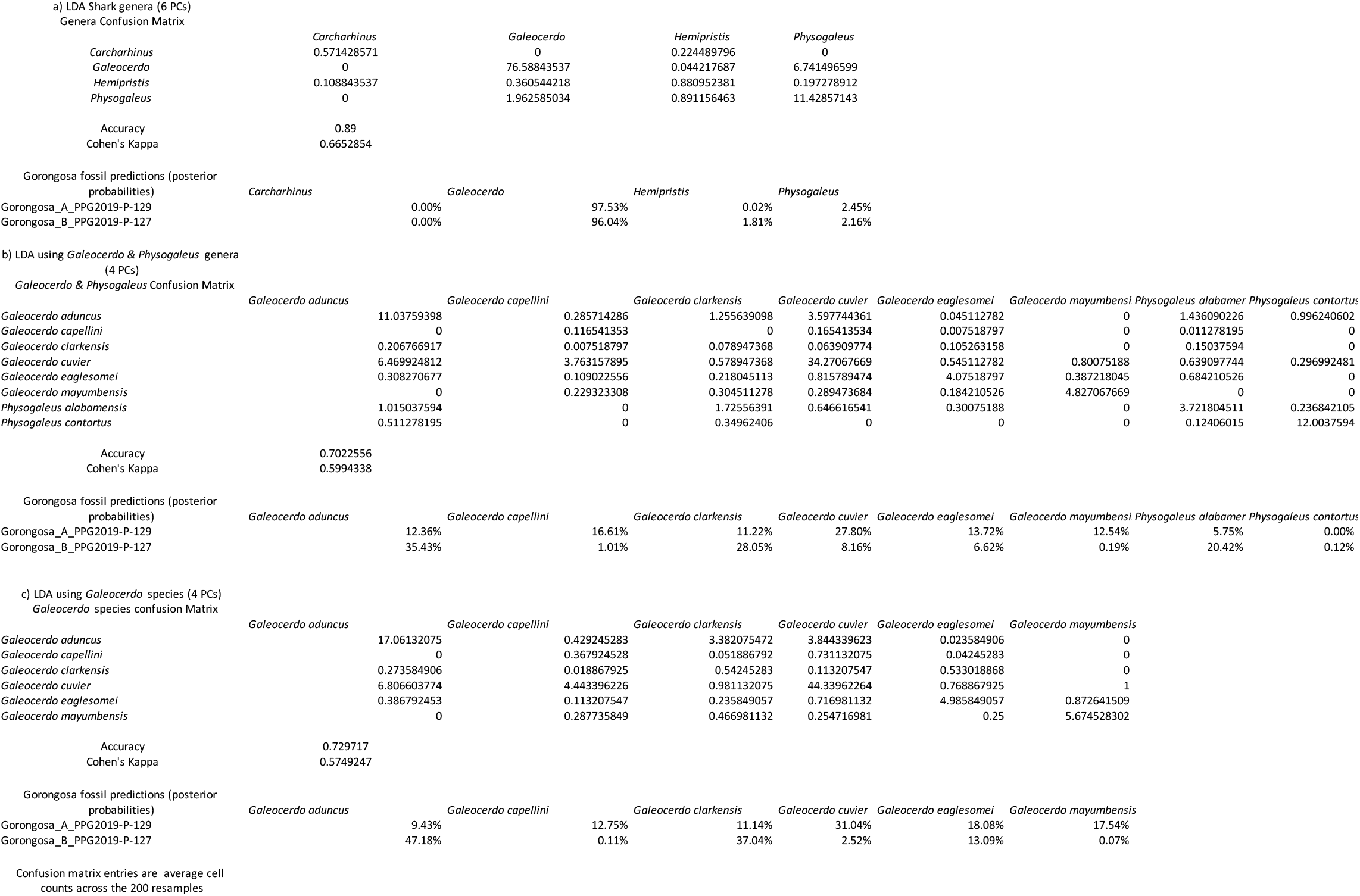
Analysis of shark teeth: results of LDA models.

**Supplementary Table S7:**
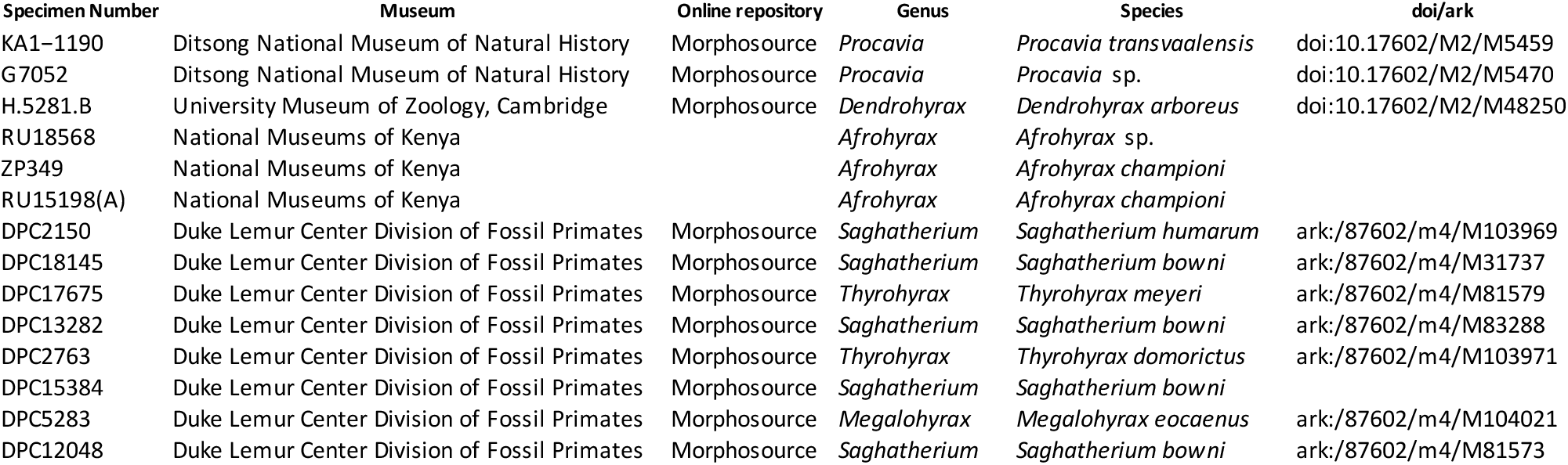
Comparative sample of hyracoid mandibular teeth from the National Museums of Kenya, Nairobi.

**Supplementary Table S8:** Comparative sample of hyracoid m3s from the National Museums of Kenya, Nairobi.

| Specimen Number | Museum | Online repository | Genus | Species | doi/ark | notes |
| --- | --- | --- | --- | --- | --- | --- |
| ZP1508 | National Museums of Kenya |  | *Bunohyrax* | *Bunohyrax* aff. *fajumensis* |  | cast |
| RU15198(A) | National Museums of Kenya |  | *Afrohyrax* | *Afrohyrax championi* |  |  |
| DPC7369 | Duke Lemur Center Division of F | Morphosource | *Thyrohyrax* | *Thyrohyrax domorictus* | ark:/87602/m4/M104159 |  |
| RU18568 | National Museums of Kenya |  | *Afrohyrax* | *Afrohyrax* sp. |  |  |
| ZP349 | National Museums of Kenya |  | *Afrohyrax* | *Afrohyrax championi* |  | cast |
| ZP347 | National Museums of Kenya |  | *Afrohyrax* | *Afrohyrax championi* |  | cast |
| ZP1211 | National Museums of Kenya |  | *Thyrohyrax* | *Thyrohyrax domorictus* |  | cast |
| WK18206(A) | National Museums of Kenya |  | *Afrohyrax* | *Afrohyrax championi* |  |  |
| DPC2763 | Duke Lemur Center Division of F | Morphosource | *Thyrohyrax* | *Thyrohyrax domorictus* | ark:/87602/m4/M103971 |  |
| DPC18145 | Duke Lemur Center Division of F | Morphosource | *Sagatherium* | *Sagatherium boweni* | ark:/87602/m4/M31737 |  |
| DPC2150 | Duke Lemur Center Division of F | Morphosource | *Sagatherium* | *Sagatherium humarum* | ark:/87602/m4/M103969 |  |
| DPC5283 | Duke Lemur Center Division of F | Morphosource | *Megalohyrax* | *Megalohyrax eoacenus* | ark:/87602/m4/M104021 |  |
| DPC12048 | Duke Lemur Center Division of F | Morphosource | *Sagatherium* | *Sagatherium boweni* | ark:/87602/m4/M81573 |  |
| NW22558 (C) | National Museums of Kenya |  | *Meroehyrax* | *Meroehyrax kyongoi* |  |  |
| DPC17675 | Duke Lemur Center Division of F | Morphosource | *Thyrohyrax* | *Thyrohyrax meyeri* | ark:/87602/m4/M81579 |  |
| DPC15384 | Duke Lemur Center Division of F | Morphosource | *Sagatherium* | *Sagatherium boweni* |  |  |
| DPC13282 | Duke Lemur Center Division of F | Morphosource | *Sagatherium* | *Sagatherium boweni* | ark:/87602/m4/M83288 |  |
| ZP1255 | National Museums of Kenya |  | *Parapliohyrax* | *Parapliohyrax mirabilis* |  | cast |
| BN802 (H) | National Museums of Kenya |  | *Parapliohyrax* | *Parapliohyrax ngororaensis* |  |  |
| LP22529 | National Museums of Kenya |  | *Thyrohyrax* | *Thyrohyrax microdon* |  |  |
| KA1-1190 | Ditsong National Museum of Na | Morphosource | *Procavia* | *Procavia transvaalensis* | doi:10.17602/M2/M5459 |  |
| G7052 | Ditsong National Museum of Na | Morphosource | *Procavia* | *Procavia* sp. | doi:10.17602/M2/M5470 |  |
| H.5281.B | University Museum of Zoology, U | Morphosource | *Dendrohyrax* | *Dendrohyrax arboreus* | doi:10.17602/M2/M48250 |  |
| NK41304 | National Museums of Kenya |  | *Dendrohyrax* | *Dendrohyrax* cf. *validus* |  |  |
| NK36934 | National Museums of Kenya |  | *Dendrohyrax* | *Dendrohyrax* cf. *validus* |  |  |

**Supplementary Table S9:** primate M1 roots comparative sample.

